# Tissue mechanics and systemic signaling safeguard epithelial tissue against spindle misorientation

**DOI:** 10.1101/2025.10.18.682384

**Authors:** Floris Bosveld, Baptiste Tesson, Eric van Leen, Sam Amirebrahimi, Raphael Thinat, Yohanns Bellaiche

## Abstract

Multicellular organisms possess conserved safeguard mechanisms that ensure the maintenance of tissue integrity. Exploring these mechanisms has proven instrumental in understanding how tissues robustly develop and prevent tumor initiation. Here, we investigate how epithelial tissues preserve their architecture and cell number in the face of spindle mis-orientation. Spindle mis-orientation, due to the lack of spindle pulling forces, centrosomes, or mitotic rounding, can cause epithelial cells to be mispositioned within or outside the tissue, leading to significant cell loss. By inducing spindle mis-orientation in *Drosophila* epithelial tissue, we first found that acentrosomal microtubules and cell contractility prevent excessive epithelial cell loss by enabling mispositioned cells to reintegrate into the epithelium. However, this mechanism alone is insufficient to maintain the total epithelial cell number. We uncovered that epithelial mechanics and cell size sensing monitor and compensate for epithelial cell loss predominantly by reducing physiological apoptosis through Hippo/YAP signaling. Lastly, we found that systemic TNF signaling protects the organism by eliminating potentially harmful non-reintegrating cells. Overall, our results delineate the complementary roles of mechanics and systemic signaling in controlling cell number and position at both tissue and organismal levels.

## Introduction

During tissue development, homeostasis, and repair, the tight control of both cell number and position is instrumental for tissue organization, stem cell dynamics, and the prevention of tumor initiation. Accordingly, tissues have implemented safeguarding mechanisms to monitor and correct defects in cell positioning or number to ensure robust tissue dynamics. Within epithelial tissues, defects in spindle orientation drastically challenge both cell positioning and number, and can promote tumor initiation.^1–7^ Yet, we lack a comprehensive understanding of the mechanisms buffering the adverse effects of epithelial cell mispositioning and loss due to spindle mis-orientation.

In monolayered epithelial tissues, cells divide with their spindles oriented in the plane of the tissue, so that both daughters are born within the tissue; thus, increasing cell number and maintaining tissue architecture.^8–10^ Epithelial spindle orientation depends on cell geometry, polarity, adhesion, mechanical, and signaling cues that converge on the cortical localization of the NuMA protein. ^8,10,11^ NuMA, in turn, binds to the Dynein complex to generate astral microtubule (MT) pulling forces that orient the spindle in the plane of the tissue.^8,10,11^ Consequently, the loss of NuMA function leads to misplaced epithelial daughter cells due to mis-oriented cell division. These misplaced cells can either reintegrate into the epithelial tissue or delaminate from it.^6,7,12–14^ While cell-cell adhesion in the *Drosophila* follicular epithelium is necessary for the reintegration of all misplaced cells within the tissue,^13,14^ in most tissues, the mechanisms regulating the balance between reintegration and delamination remain unknown. Importantly, upon spindle mis-orientation, daughter cells failing to reintegrate into the tissue undergo apoptosis, resulting in cell divisions that produce only a lone epithelial resident daughter.^6,7^ Despite this reduced increase in cell number upon division mis-orientation, the loss of NuMA has been reported to cause only mild effects on tissue development,^7,12^ suggesting the existence of mechanisms compensating for cell divisions producing only one epithelial daughter cell. In addition, the death of delaminated cells is shown to be essential to avoid tumor initiation.^8^ However, the mechanisms triggering their apoptosis are still unknown. Notably, both the loss of centrosomes and the lack of mitotic cell rounding also cause spindle orientation defects.^7,15–19^ This further emphasizes the need to comprehensively characterize the mechanisms that promote cell reintegration, compensate for epithelial cell loss, and ensure the death of delaminated daughter cells.

Here, we investigate the mechanisms controlling cell positioning and number upon spindle mis-orientation by taking advantage of high-resolution and long-term imaging in the *Drosophila* pupal dorsal thorax epithelium (notum) within a whole living organism. Using *Drosophila* epithelial tissue devoid of the conserved spindle guidance protein Mud (mammalian NuMA), we first delineate the mechanisms controlling the balance between reintegration and delamination by uncovering that acentrosomal MTs and cortical tension are necessary for cell reintegration upon division mis-orientation. Second, we found that upon the delamination of a misplaced daughter cell, the apex of the lone epithelial daughter served as a mechanical ruler for cell number control, reducing the apoptosis rate in the tissue and thus, restoring tissue cell number. Lastly, we established that systemic TNF signaling promotes the death of delaminated cells. Altogether, our results reveal how the complementary roles of cell mechanics and systemic signaling act as safeguard mechanisms in the control of cell positioning and number at the tissue and animal level.

## Results

### Spindle mis-orientation leads to epithelial cell mispositioning and loss

We have previously reported that *Drosophila* NuMA, Mud, controls the apical-basal (AB) spindle orientation in the posterior domain of the *Drosophila* pupal notum, the scutellum.^20,21^ Therefore, we decided to focus on this region to explore the short- and long-term consequences of spindle mis-orientation on epithelial cell dynamics.

Within notum tissues expressing the E-Cad:GFP apical adherens junction (AJ) marker, cell divisions can be easily identified by cell apex area expansion and rounding during mitosis.^20,21^ Upon division, we thus tracked and compared the behaviors of the two daughter cells in wild-type (wt) and *mud* scutellum tissues (Figure 1A; Video S1). As expected, in wt tissue the two daughter cells are positioned within the tissue, and their apical areas are of similar size. In *mud* tissue, 60% of the divisions (*n=* 4829 out of 7975 divisions) initially produced 2 daughter cells, both of which can be easily distinguished within the tissue upon cytokinesis. The initial size of the two daughter apices varied from nearly equal to differences as high as 90%; despite such difference in their initial apex size the two daughters rapidly reached similar apex size (Figure 1A; Video S1). The remaining 40% of the cell divisions can be further categorized as follows. In 27% of the cases (12% of the total number of division), cell division leads to the formation of one very large daughter cell and a barely identifiable second daughter (apex area less than 10% of its sibling). The apex of this second daughter increased in area as the cell reintegrated within the tissue, eventually leading to the presence of two daughter cells with approximately equal apex area. In the remaining 73% of cases (27% of the total number of division), only one daughter was observed in the tissue with a disproportionally large apex area (Figures 1A and 1B; Video S1). Time-lapse imaging using E-Cad:GFP, the nuclear His2B:RFP and centrosomal YFP:Asl markers in *mud* tissue showed that such mis-oriented division led to one sibling located below the epithelial tissue (Figure 1C). These basal daughter cells eventually underwent apoptosis as shown by the accumulation of cleaved Caspase-3 (Casp3) positive staining in cells beneath *mud* epithelial tissue at 22 hours after pupa formation (hAPF) (Figures 1D-F). These later divisions therefore produce a lone epithelial daughter, leading to substantial epithelial cell loss. Importantly, the formation of one or two epithelial daughter cells upon cell division within *mud* tissue correlated with the magnitude of AB spindle mis-orientation (Figure 1G). As found in multiple tissues,^6,7,12–14,22,23^ spindle mis-orientation can cause both epithelial cell mispositioning and cell loss in the notum. To investigate how cell positioning and number are regulated upon division mis-orientation, we focused on the 40% of divisions with a highly tilted spindle, which produced either two epithelial daughters, one of which has an apex area less than 10% of its sibling, or a single lone epithelial daughter cell.

**Figure 1.**
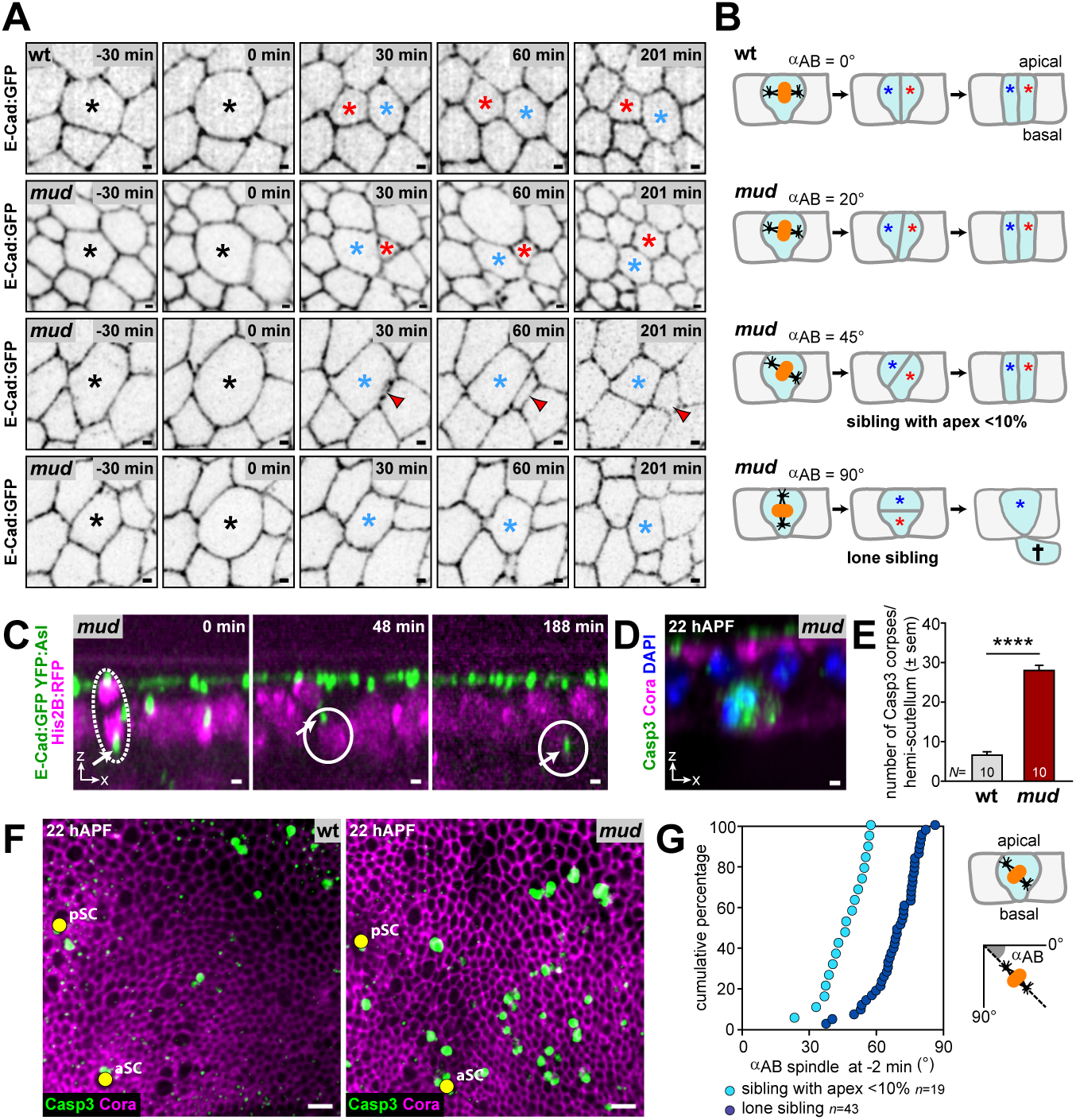
Division mis-orientation induces cell mispositioning and loss. (A) Top view time-lapse images of apical E-Cad:GFP in wt (top panels) and *mud* tissues. wt cells divide with an average apex area difference of 14.55 ± 1.75 sem (*n=* 50 cells). For *mud* three types of divisions are shown. Divisions that produce two daughter cells, one whereby both daughters have a 60% difference in apex area (second row panels), and a division where one daughter cell has an apex area <10% of its sibling (third row panels), as well as a division that produces a lone epithelia daughter cell (bottom panels). Black asterisks, dividing cells at anaphase onset *t=* 0 min. Red and blue asterisks indicate tissue resident daughter cells following division characterized by a small or large apex, respectively. Red arrowheads, reintegrating mispositioned daughter cell. (B) Apical-basal (AB) schematics illustrating the orientation of the spindle and the positioning of daughter cells in wt and *mud* mutant tissues. The relative apical areas of the two daughter cells for each division type are indicated and correspond to divisions shown in A. The orientation (°) of the mitotic spindle along the apical-basal axis (αAB) is indicated for each panel. (C) AB view time-lapse images of E-Cad:GFP, YFP:Asl and His2B:RFP in *mud* tissue showing a highly mis-oriented division (dashed white outline) resulting in a basally born cell residing underneath the epithelium (solid white outlines). White arrows, centrosome of the basally born daughter cell. (D) AB view image of fixed *mud* tissue at 22 hAPF stained with antibodies against SJ marker Cora and cleaved Casp3 showing a basal apoptotic Casp3 positive corpse. (E) Graph of the number (mean ± sem) of Casp3 positive corpses per hemi-scutellum tissue in wt and *mud*. **** *p*<0.0001, student *t*-test. *N*, number of tissues analyzed. (F) Top view images of fixed wt and *mud* tissues at 22 hAPF stained with antibodies against SJ marker Cora and cleaved Casp3. aSC and pSC, anterior and posterior scutullar macrochaetes (yellow circles). (G) Graph of αAB at *t=* -2 min prior to anaphase onset for *mud* divisions that produced either two epithelial daughters whereby one cell has and apex area <10% of its sibling (light blue) or a lone epithelial daughter (dark blue), plotted as cumulative percentage. The two distributions are significantly different (*p*<0.0001, Mann-Whitney U test). Schematic, αAB was calculated by measuring the spindle orientation relative to the tissue AB axis, whereby a fully planar orientation is characterized by an angle of 0°, while a perpendicular orientation by an angle of 90° (see B). *n*, number of divisions analyzed. Scale bars: 1 µm (A, C, D), 10 µm (F).

### Lateral cell-cell adhesion does not modulate basal-to-apical reintegration of mispositioned cells

We first aimed to understand the mechanisms that ensure the reintegration of highly mispositioned daughter cells, thereby preventing their loss. Upon spindle mis-orientation in the *Drosophila* follicular epithelium, which results solely in apical cell mispositioning, all apically misplaced cells reintegrate back into the epithelium. In this tissue with immature septate junctions (SJ), such apical-to-basal reintegration depends on lateral cell-cell adhesion SJ proteins, Nrg, FasII and FasIII.^13,14,24^ The notum tissue harbors mature SJ^25,26^ and as observed in other tissues, basally mispositioned cells reintegrate within the tissue in a basal-to-apical reintegration process.^12,13^ To test whether basal-to-apical reintegration depends on lateral adhesion, we knocked-down Nrg, FasII or FasIII function by RNAi in clones generated within *mud* tissues. The ratio of delaminating over reintegrating cells (R_d/r_) were then compared between the double loss-of-function clones and surrounding *mud* tissue. The R_d/r_ ratios were similar in *mud* and in *mud* tissue upon knock-down of *Nrg^RNAi^*, *FasII^RNAi^* or *FasIII^RNAi^* (Figures 2A and S1A-C) indicating that lateral adhesion is not essential for basal-to-apical cell reintegration.

**Figure 2.**
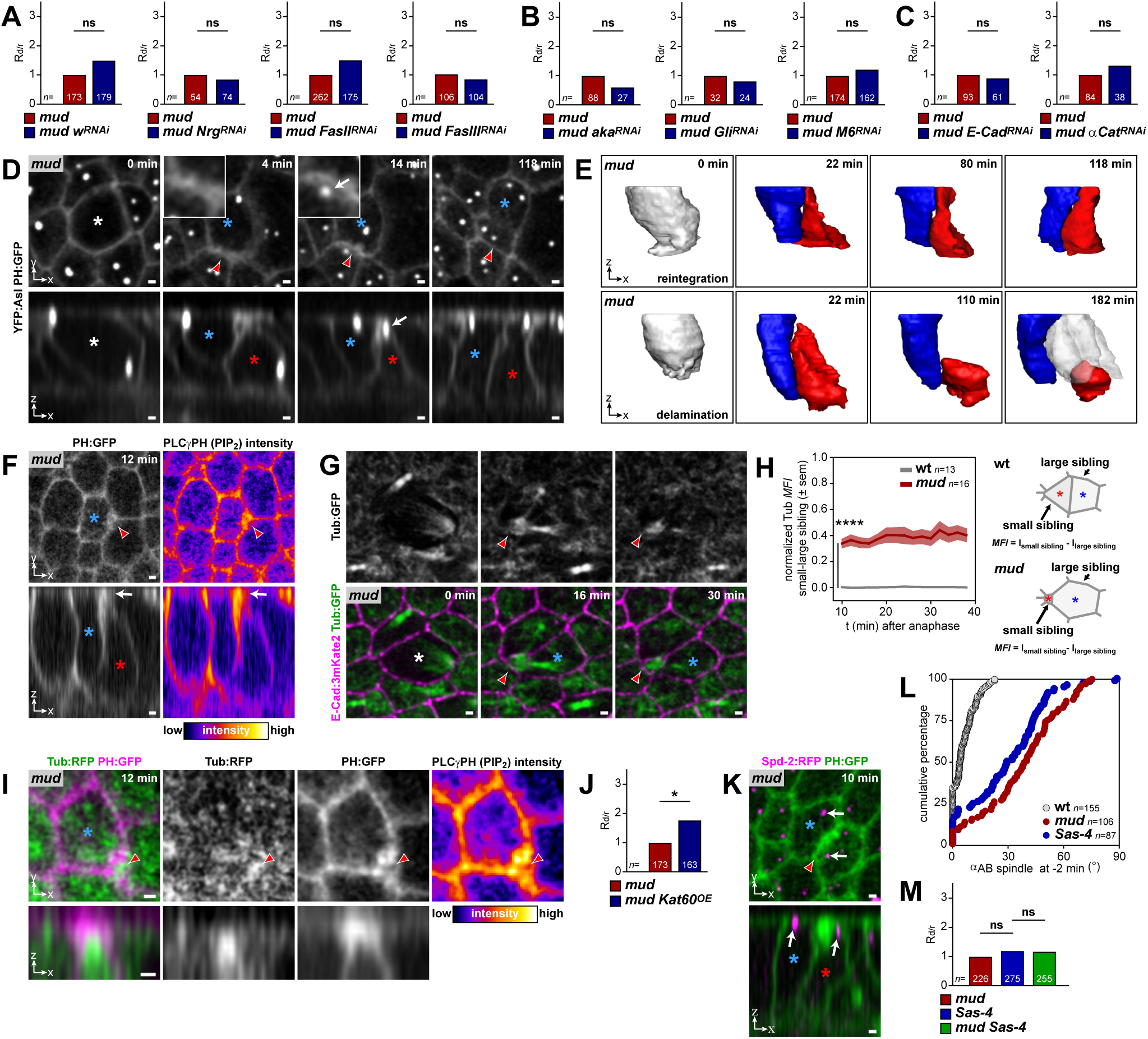
Cell reintegration is associated with the formation of an apical cap enriched in PIP_2_ and microtubules. (A) Graphs of the delamination/reintegration ratios (R_d/r_) in *mud*, and *mud w^RNAi^*, *mud Nrg^RNAi^*, *mud FasII^RNAi^*, *mud FasIII^RNAi^* tissue normalized to the R_d/r_ in *mud*. ns, not significant, chi-square test. *n*, number of divisions analyzed. (B) Graphs of the delamination/reintegration ratios (R_d/r_) in *mud*, and *mud aka^RNAi^*, *mud Gli^RNAi^*, *mud M6^RNAi^* tissue normalized to the R_d/r_ in *mud*. ns, not significant, chi-square test. *n*, number of divisions analyzed. (C) Graphs of the delamination/reintegration ratios (R_d/r_) in *mud*, and *mud E-cad^RNAi^*, *mud αCat^RNAi^* tissue normalized to the R_d/r_ in *mud*. ns, not significant, chi-square test. *n*, number of divisions analyzed. (D) Top (top panels) and AB (bottom panels) view time-lapse images of YFP:Asl and PH:GFP of a highly mis-oriented division in *mud* tissue resulting in a mispositioned daughter cell (red arrowheads in top views and red asterisk in AB views) that reintegrates back within the tissue. Insets show high magnification of the apical cap formed by the reintegrating cell. Note that the centrosome appears in the cap at *t=* 14 min (white arrows). White asterisk, dividing cell at anaphase onset *t=* 0 min. Blue asterisk, large sibling. (E) 3D segmented time-lapse images (AB views) of a mis-oriented division resulting in a mispositioned daughter cell that reintegrates back within the tissue (top panels, same cell as shown in D), or a lone epithelial daughter cell (bottom panels) captured from *mud* epithelia expressing YFP:Asl and PH:GFP. The blue cells correspond to either the large sibling or the lone epithelial daughter resulting from a mis-oriented division, while the red cells correspond to the reintegrating or delaminating siblings, respectively. (F) Top (top panels) and AB (bottom panels) view live images of PH:GFP (right panels) and the intensity of the PLCγPH(PIP2) reporter in Fire LUT (right panels) at *t=* 12 min after anaphase onset. Red arrowheads, reintegrating cell. Blue asterisk, large sibling. Red asterisk, reintegrating daughter. White arrows, apical cap (corresponding to the reintegrating cell, red arrowheads). (G) Top view time-lapse images of apical Tub:GFP (grey top panels, magenta bottom panels) and E-Cad:3mKate2 (magenta bottom panels) in *mud* tissue showing a highly mis-oriented division resulting in a mispositioned daughter cell that reintegrates back within the epithelium (red arrows). White asterisk, dividing cell at anaphase onset *t=* 0 min. Blue asterisks, large sibling. (H) Graph of the evolution of the normalized Tub:GFP mean apical fluorescence intensity (*MFI* ± sem) measured in the smallest daughter cells as compared to their larger siblings upon division in wt and *mud* tissues. **** *p*<0.0001, student *t*-test at *t=* 10 min. *n*, number of cells analyzed. The schematic to the right shows how the normalized *MFI* was calculated in *mud* and wt tissue. Thereto the apical intensity (I) in the largest sibling was subtracted from the intensity in the smallest daughter cell (in *mud* this corresponds to reintegrating cells with an initial apex area <10% of its sibling, while in wt this reflects the daughter cells with the smallest apex area). The resulting intensities were normalized to the highest measured difference (see Methods). Asterisks indicate tissue resident daughter cells characterized by a small (red) or large (blue) apex following division in wt and *mud* tissue. (I) Top (top panels) and AB (bottom panels) view live images of the apical cap (red arrowheads) localization of Tub:GFP and PH:ChFP at *t=* 12 min after anaphase onset in a reintegrating cell in *mud* tissue. The PLCγPH(PIP_2_) intensity is shown in Fire LUT. Blue asterisk, large sibling. (J) Graph of the delamination/reintegration ratio (R_d/r_) in *mud*, and *mud Kat60^OE^* tissue normalized to the R_d/r_ in *mud*. * *p*<0.05, chi-square test. *n*, number of divisions analyzed. (K) Top (top) and AB (bottom) view live images of PH:GFP and Spd-2:RFP in *mud* tissue following a highly mis-oriented division at *t=* 10 min after anaphase onset. Blue asterisk, large sibling. Red arrowhead/asterisk, reintegrating cell. White arrows, centrosomes. (L) Graph of áAB at *t=* -2 min prior to anaphase onset in wt, *mud* and *Sas-4*, plotted as cumulative percentage. The αAB in *mud* and *Sas-4* are significantly different from wt (*p*<0.0001, Mann-Whitney U test). *n*, number of cells analyzed. (M) Graph of the delamination/reintegration ratio (R_d/r_) in *mud*, *Sas-4*, and *mud Sas-4* tissue normalized to the R_d/r_ in *mud*. ns, not significant, chi-square test. *n*, number of divisions analyzed. Scale bars: 1 µm.

We then tested whether other regulators of cell-cell adhesion might promote cell reintegration. Basal-to-apical cell insertion, or radial cell intercalation, occurs as part of mucociliary epitheliogenesis during *Xenopus* embryogenesis.^27,28^ Since during this process, tricellular junctions (TCJ) and their components play critical roles,^28^ we investigated whether known *Drosophila* lateral tricellular SJ components Aka^29^, Gli^30^ and M6^31^ would facilitate the basal-to-apical reintegration. As for the depletions of the lateral adhesion proteins, no changes in the R_d/r_ ratios were observed when comparing *mud* tissues with surrounding *mud aka^RNAi^*, *mud Gli^RNAi^* or *mud M6^RNAi^* clones (Figures 2B and S1D-F). These results indicate that tricellular SJ components are unlikely to ensure basal-to-apical reintegration of the mispositioned cells. Lastly, we tested the role of the core AJ cell-cell adhesion components E-Cad and *α*-Catenin.^32,33^ Neither *E-Cad^RNAi^* nor *αCat^RNAi^* knock-down in *mud* tissues modified the R_d/r_ upon spindle mis-orientation (Figures 2C, S1G and S1H). Based on these results, we conclude that the reintegration of basally mispositioned cells is not affected by decreasing the function of components of the lateral and tricellular septate junctions, or adherens junctions.

### The initial step of cell reintegration is controlled by acentrosomal microtubules

To understand how mispositioned cells reintegrate within the tissue in a basal-to-apical process, we then characterized the 3D shape of reintegrating cells versus delaminating cells upon spindle mis-orientation. Using the membrane marker PLCγPH:GFP (PH:GFP), which binds phosphatidylinositol biphosphate (PIP_2_),^34^ to record the 3D cell shape and YFP:Asl to follow spindle orientation, we analyzed *mud* daughter cells generated by a division with a spindle tilt greater than 40°, since many of them either reintegrate or delaminate from the tissue (Figure 1G). We found that most of the daughters that reintegrated within the tissue initially formed an apical cap (91%, *n=* 69 cells), whereas 98% of daughter cells without an apical cap delaminated from the tissue (*n=* 107 cells, Figure 2D). This apical cap constitutes an apical membranous structure enriched in PIP_2_ (Figure 2F). 3D cell segmentation revealed that the apical cap belongs to an elongated apical-to-basal stalk formed by the mispositioned daughter (Figure 2E). The cap then progressively enlarged as the mispositioned daughter cell reintegrated in the tissue (Figures 2D and 2E; Video S2). This suggests that the initial step for cell reintegration is associated with the formation and maintenance of a membranous apical cap.

To define how this initial step of cell reintegration is regulated, we aimed to identify proteins enriched in the apical cap. Strikingly, during our analyses of spindle orientation using the MT marker *α−*Tubulin (Tub:GFP), we observed an enrichment of MTs in the apical cap of the reintegrating daughter cells as compared to their larger siblings (Figures 2G-I; Video S3), implicating MTs as potential regulators of cell reintegration. Indeed, over-expression of the MT severing enzyme Kat60^35,36^ in *mud* tissue significantly increased the R_d/r_ (Figure 2J), while did not elicit a phenotype in wt interphasic or mitotic cells (*n*= 149 cells). Thus, MTs contribute to the reintegration of highly mispositioned cells within the tissue to prevent their loss.

MTs can be nucleated by centrosomal or non-centrosomal mechanisms.^16,37^ Since we found that centrosomes were present in the apical cap (Figures 2D and 2K) and centrioles have been reported to control radial cell intercalation in *Xenopus*,^38^ we first investigated the putative role of centrosomes. This was aided by analyzing divisions in *Sas-4* tissues, which are devoid of centrosomes and known to cause spindle mis-orientation in epithelial tissues.^7,16,39^ In *Sas-4* notum tissues, we found that mitotic spindles were mis-oriented resulting in cell mispositioning as observed in *mud* (Figures 2L and 2M). We found that 11% (*n=* 613 out of 5445 divisions) of the of divisions led to the formation of one barely visible daughter cell (apex area <10% of its sibling) that subsequently reintegrated within the tissue. This number is close to that in *mud* mutant tissue, Furthermore, the R_d/r_ was similar in *mud*, *Sas-4* and *mud Sas-4* double mutant tissues (Figure 2M), indicating that centrosomes do not contribute to the reintegration of basally mispositioned *mud* cells. Together, these results prompted us to investigate whether regulators of non-centrosomal MTs might contribute to cell reintegration.

The MT minus-end stabilizing protein Patronin (Pat, CAMSAP) has important functions in the regulation of non-centrosomal MT dynamics.^40–44^ Interestingly, a functional Pat:tagRFP^45^, whose localization is restricted to the apical domain in notum tissue (Figures S1I and S1J), localizes to the apical cap and is enriched in the apex of the reintegrating cells during reintegration as compared to their larger siblings (Figures 3A and 3B; Video S3). Importantly, no enrichment in apical Pat:tagRFP (*n=* 8 cells) or Tub:RFP (*n=* 9 cells) was observed in mispositioned cells that did not form an apical cap (Figures S1K and S1L). Furthermore, while over-expression of Pat did not elicit a phenotype in *mud* tissue, we found that the depletion of Pat by clonal dsRNA expression in *mud* tissue increased the R_d/r_ ratio by 2.6-fold (Figures 3C, S1M and S1N). We then investigated the specificity of the Pat loss-of-function regarding cell reintegration. dsRNA knock-down of Pat function neither led to cell delamination in wt tissue, nor affected spindle orientation in wt or *mud* tissue, nor changed the 3D cell geometry compared to that observed in *mud* tissue (Figures S1O-R), suggesting a specific role during cell reintegration. Consistent with an acentrosomal MT-dependent mechanism preventing cell loss, Pat localization and dynamics during reintegration were similar in reintegrating *mud* and *Sas-4* cells (Figures S1S and S1T). Together, our data suggest that the acentrosomal function of Pat in MT dynamics or organization promotes the reintegration of highly mispositioned daughter cells.

**Figure 3.**
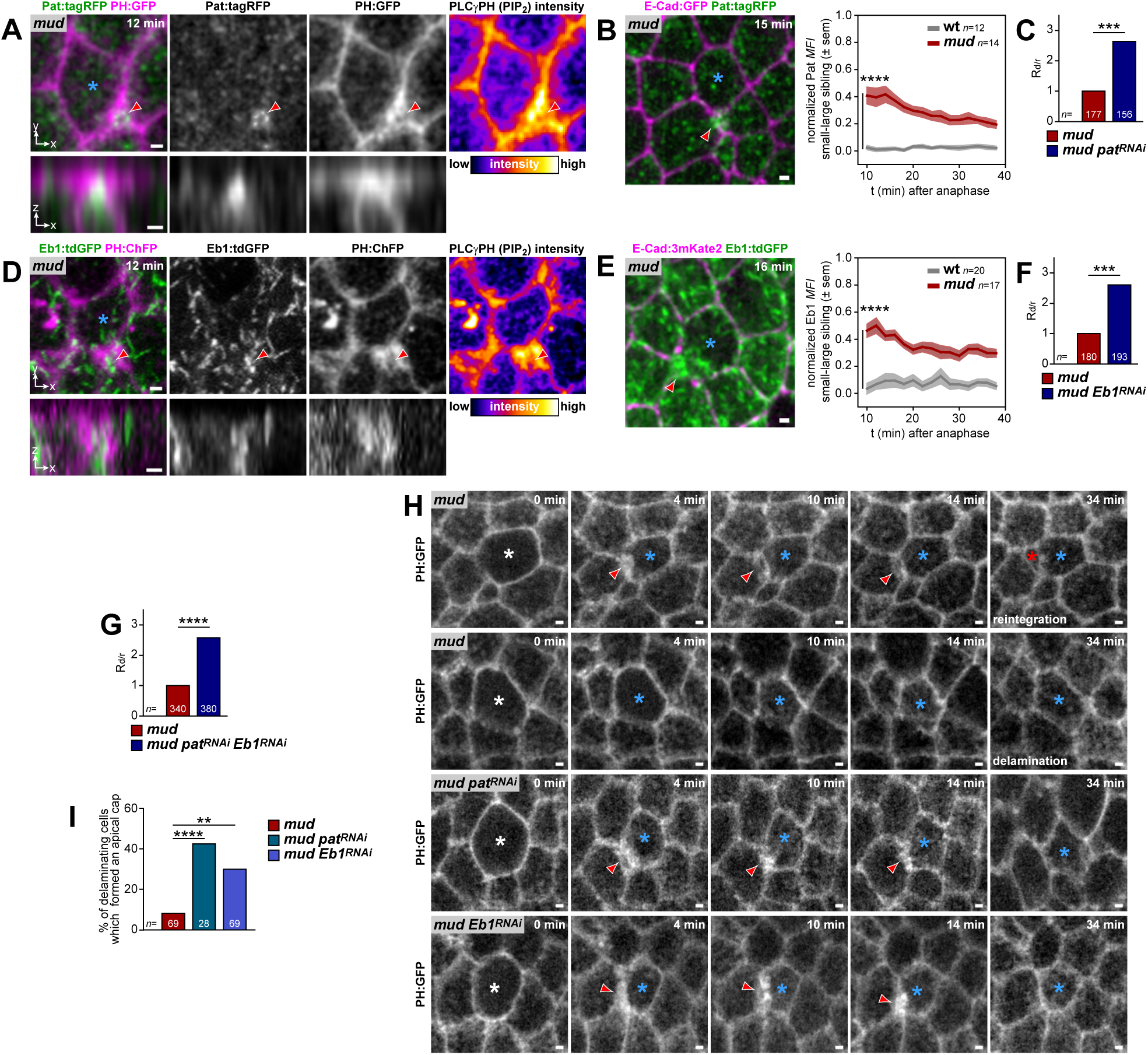
Pat/Eb1 are necessary to initiate the reintegration of mispositioned cells. (A) Top (top panels) and AB (bottom panels) view live images of the apical cap (red arrowheads) localization of Pat:tagRFP and PH:GFP at *t=* 12 min after anaphase onset in a reintegrating cell in *mud* tissue. The PLCγPH(PIP_2_) intensity is shown in Fire LUT. Blue asterisk, large sibling. (B) The left top view live image shows apical Pat:tagRFP localization at *t=* 15 min after anaphase onset in a small daughter cell (red arrowhead) following a highly mis-oriented division in *mud* tissue co-expressing E-Cad:GFP. Blue asterisk, large sibling. The right graph shows the evolution of the normalized Pat:tagRFP mean apical fluorescence intensity (*MFI* ± sem) measured in the smallest daughter cells as compared to their larger siblings upon division in wt and *mud* tissues. **** *p*<0.0001, student *t*-test at *t=* 10 min. *n*, number of cells analyzed. (C) Graph of the delamination/reintegration ratio (R_d/r_) in *mud*, and *mud pat^RNAi^* tissue normalized to the R_d/r_ in *mud*. *** *p*<0.001, chi-square test. *n*, number of divisions analyzed. (D) Top (top panels) and AB (bottom panels) view live images of the apical cap (red arrowheads) localization of Eb1:tdGFP and PH:ChFP at *t=* 12 min after anaphase onset in a reintegrating cell in *mud* tissue. The PLCγPH(PIP_2_) intensity is shown in Fire LUT. Blue asterisk, large sibling. (E) The left top view live image shows apical Eb1:tdGFP localization at *t=* 16 min after anaphase onset in a small daughter cell (red arrowhead) following a highly mis-oriented division in *mud* tissue co-expressing E-Cad:3mKate2. Blue asterisk, large sibling. The right graph shows the evolution of the normalized Eb1:tdGFP mean apical fluorescence intensity (*MFI* ± sem) measured in the smallest daughter cells as compared to their larger siblings upon division in wt and *mud* tissues. **** *p*<0.0001, student *t*-test at *t=* 10 min. *n*, number of divisions. (F) Graph of the delamination/reintegration ratio (R_d/r_) in *mud*, and *mud Eb1^RNAi^* tissue normalized to the R_d/r_ in *mud*. *** *p*<0.001, chi-square test. *n*, number of divisions analyzed. (G) Graph of the delamination/reintegration ratio (R_d/r_) in *mud*, and *mud pat^RNAi^ Eb1^RNAi^* tissue normalized to the R_d/r_ in *mud*. **** *p*<0.0001, chi-square test. *n*, number of divisions analyzed. (H) Top view time-lapse images of apical PH:GFP localization in *mud*, *mud pat^RNAi^* and *mud Eb1^RNAi^*. Upon a highly mis-oriented division in *mud* tissue, most cells (91%, *n=* 69 cells) forming an apical cap (red arrowheads) reintegrated within the epithelium (red asterisk, top panels). On the contrary, such apical cap was mostly absent in divisions that produced a delaminating daughter cell (blue asterisks, 5% of *n=* 111 delaminated cells had a cap) (second row panels). In *mud pat^RNAi^* and *mud Eb1^RNAi^* (bottom two panels) mis-oriented divisions associated with the formation of an apical cap (red arrowheads) frequently produced a lone epithelial daughter (blue asterisks, quantified in I). White asterisks, dividing cells at anaphase onset *t=* 0 min. Blue asterisks, large or lone siblings. (I) Graph of the percentage of mis-oriented divisions that produced daughters forming an apical cap and that subsequently delaminated from the epithelium in *mud*, *mud pat^RNAi^* and *mud Eb1^RNAi^*. The frequency of cells forming an apical cap and delaminated rather than reintegrated back within the tissue was significantly different between *mud* and *mud pat^RNAi^* (*p*<0.0013) or *mud* and *mud Eb1^RNAi^* (*p*<0.0000). ** *p*<0.01, **** *p*<0.0001, chi-square test. *n*, number of divisions analyzed. Scale bars: 1 µm.

To analyze whether Pat localization at in the apex of the reintegrating cell is relevant for cell reintegration, we aimed the define the mechanism by which Pat is apically localized within the apex of reintegrating cells. To this end we screened through a set of 16 genes, known to interact with Pat or reported to be enriched at the apical side of epithelial cells (Figures S2A-K).^42,46–57^ For these selected candidates, we analyzed in *mud* tissue either both their localization in the apical cap during cell reintegration as well as their effect on the R_d/r_ or only their effect on the R_d/r_ (Figures S2A-K). This screen revealed that the apical cap was enriched in several epithelial apical components including the Sas transmembrane protein (Figure S2J). Sas encodes an apical surface protein involved the elimination of tumorigenic cells.^58–60^ Furthermore, we observed that the depletion of Sas by clonal dsRNA expression in *mud* tissue specifically increased the R_d/r_ ratio (Figure S2K). Accordingly, upon knockdown of Sas, Pat is displaced from the medial apical cortex of interphasic and reintegrating cells to the cell-cell junctions (Figures S1M and S1N). This suggest that medial Pat localization at the cell apex is necessary for cell reintegration and that the apical epithelial protein Sas controls the reintegration of mispositioned cells upon spindle mis-orientation.

In parallel, we explored the function of the plus-end binding MT polymerizing protein Eb1, which also plays a critical role in acentrosomal MT dynamics.^44,61^ Mirroring the results for Pat, we found that Eb1 was enriched in the apical cap of reintegrating cells and absent from daughter cells that failed to reintegrate (Figures 3D, 3E, and S3A). Furthermore, the depletion of Eb1 by RNAi in *mud* tissue increased the R_d/r_ ratio by 2.6-fold (Figures 3F and S3B). As shown for Pat, this Eb1 enhancement was specific to cell reintegration and Eb1 over-expression in *mud* tissue did not modulate the R_d/r_ ratio (Figure S3C). Last, triple *mud pat^RNAi^ Eb1^RNAi^* mutant tissues exhibited a R_d/r_ ratio similar to that of *mud pat^RNAi^* or *mud Eb1^RNAi^*, indicating that Pat and Eb1 are likely involved in the same process to ensure epithelial cell repositioning of misplaced cells upon defects in division orientation (Figure 2G).

Since cell reintegration is initiated by the formation of an apical cap in the reintegrating cells, we then analyzed its formation and dynamics in mispositioned *mud pat^RNAi^* and *mud Eb1^RNAi^* cells in tissue expressing the PH:GFP membrane marker. In contrast to *mud* tissue where most mispositioned cells forming a cap subsequently reintegrated within the tissue, we found that a large proportion of mispositioned cells forming an apical cap subsequently delaminated in *mud pat^RNAi^* and *mud Eb1^RNAi^* cells (Figures 2H and 2I; Video S4). We then compared the reintegration dynamics of mispositioned daughter cells in *mud* tissue with those in *mud Eb1^RNAi^* and *mud pat^RNAi^* tissues that eventually reintegrated within the tissue. The reintegration dynamics, as measured by the expansion of their apex areas relative to their siblings between 10 to 500 min after anaphase onset, was similar in *mud*, *mud Eb1^RNAi^*, and *mud pat^RNAi^* cells (Figures S3G and S3H). Taken together, we conclude that, in *mud* tissue, Pat and Eb1 are necessary to maintain or stabilize the apical cap, but not to form it, thereby promoting the basal-to-apical reintegration of mispositioned daughter cells and preventing their loss. Interestingly, although Pat and Eb1 apical intensity levels remained high and largely unchanged during early (10-38 min after anaphase onset) reintegration (Figures S3I-K), they are not required for subsequent apex area expansion once an apical cap has been successfully formed and stabilized.

### Local cortical tension contributes to cell reintegration

Having found that the initial step of cell reintegration is associated with the formation of an apical cap and that the MT regulators Eb1 and Pat are required for its maintenance to allow reintegration to proceed, we next aimed to identify the mechanisms driving the subsequent step of apex area expansion during reintegration. In particular, we sought to investigate the roles of the surrounding cells by analyzing whether reintegrating cells push on the surrounding cells or are being pulled by them. Towards this goal, we first performed two types of laser ablation experiments. First, we ablated the medial apical apex of the reintegrating cells. Upon ablation, the cell apical area of these cells very rapidly increased (Figures 4A and 4B; Video S5). Second, when we ablated the cells surrounding a reintegrating cell, the apical area of the isolated cell decreased by approximately 60% (Figures 4C and 4D; Video S5). Additionally, when we analyzed reintegrating cells neighboring a dividing cell undergoing cytokinesis, we observed that cytokinetic ring constriction temporally sped up the reintegration of mispositioned cells (Figures 4E-G). Since cytokinetic ring constriction produces pulling forces on neighboring cells,^62–64^ combined these findings agree with the notion that pulling forces from neighboring cells may modulate the reintegration of mispositioned cells.

MyoII being a critical regulator of junctional tension,^32,33,65,66^ we decided to analyze this possible neighboring pulling effect in more detail, first by investigating *mud* tissue in which we over-expressed a dominant negative form of MyoII heavy chain (*zip^DN^*)^67,68^ to decrease MyoII function. In *mud zip^DN^* tissue, a large portion of daughter cells that exhibited an initially small apex due to spindle mis-orientation failed to further expand their apices, which then shrank and disappeared (Figures 4H and 4I; Video S6). Moreover, for *mud zip^DN^* mispositioned daughter cells that eventually managed to reintegrate within the tissue, we found that they exhibited slower reintegration dynamics as compared to control *mud w^RNAi^* tissue (Figure 4J). To further test the contribution of the neighboring cells, we analyzed the dynamics of reintegrating *mud* cells that retain MyoII function themselves, abutting *mud* tissue in which we clonally over-expressed *zip^DN^* (*mud-mud zip^DN^*) and compared them with reintegrating *mud* cells neighboring a control *mud w^RNAi^* tissue (*mud-mud w^RNAi^*). As observed for the global inactivation of MyoII activity, the reintegration of cells was slower for cells facing neighboring cells compromised of MyoII function as compared to the control ones (Figures 4K and 4L; Video S6). Similar results were obtained when we analyzed *mud* cells facing *mud* cells in which we over-expressed a dsRNA against the positive regulator of MyoII activity, Rok^69^ (Figure 4M). This role of neighboring cells concurred with the observation that the apices of reintegrating cells exhibited lower MyoII enrichment compared to that of the neighboring cells (Figure S4A). Importantly, reporters of actomyosin activity (Rok:GFP, AniRBD:GFP, UtrABD:GFP) showed no apparent enrichment in neighboring cell junctions (Figure S4B). Accordingly, we did not observe a change in the junctional tension in cells neighboring a reintegrating cell (+1N) as compared to the rest of the tissue (+2N) in *mud* tissues (Figures S4C and S4D). Hence, reintegration of highly mispositioned cells depends on actomyosin forces, but does not require their upregulation in neighboring cells; instead, these forces originate from the tensile forces present within the epithelium. Consistent with these findings, we did not detect a change in neighboring cell junction tension or MyoII intensity in *mud w^RNAi^* or *mud pat^RNAi^* double mutant tissue (Figures S4E and S4F). Combined with the observation that the reintegration dynamics of *mud Eb1^RNAi^* or *mud pat^RNAi^* double mutant cells were similar to *mud w^RNAi^* (Figures S3G and S3H), these findings show that MyoII-dependent pulling forces in neighboring cells are likely independent of acentrosomal MT regulation in reintegrating cells.

**Figure 4.**
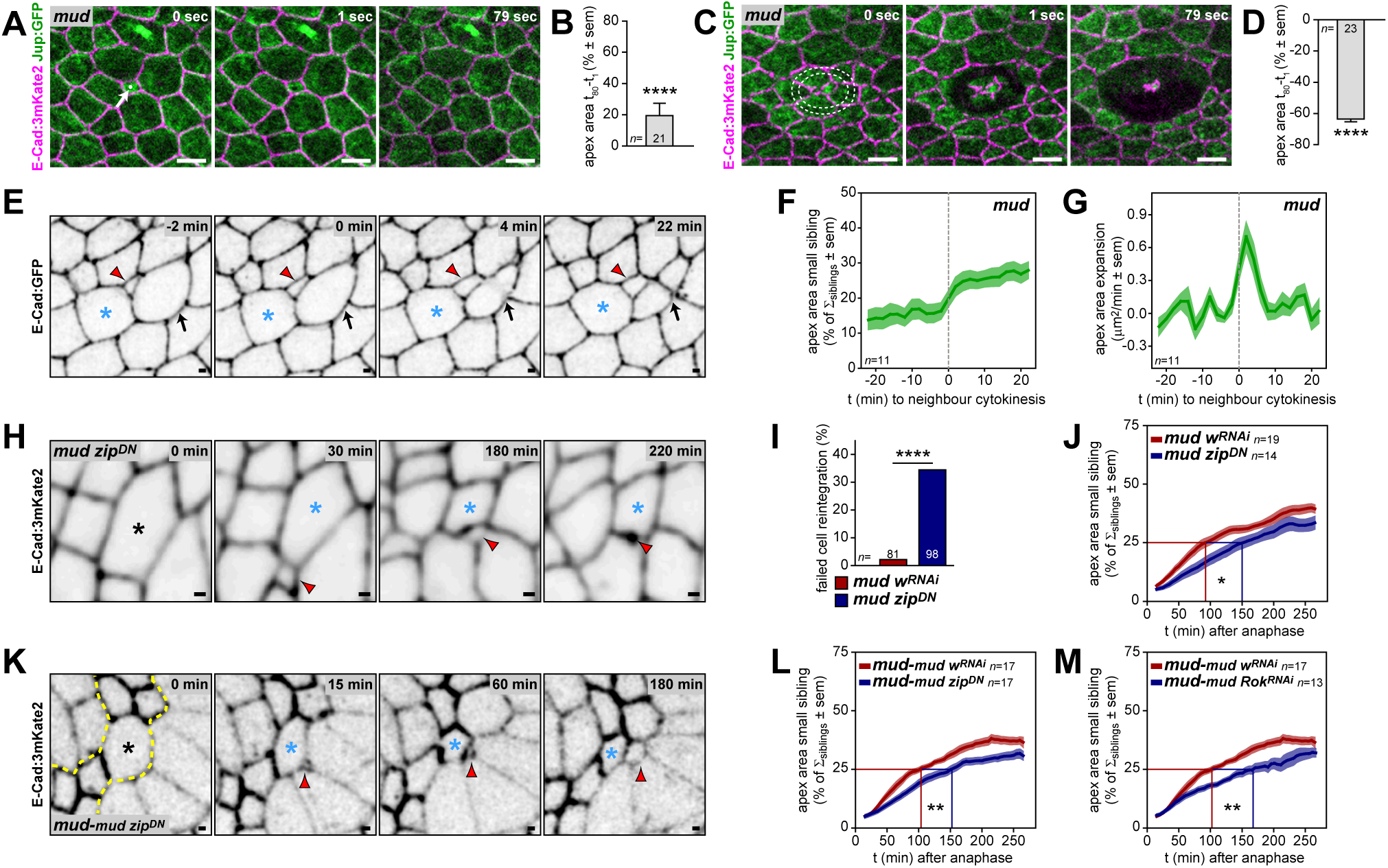
Tissue tension promotes reintegration of mispositioned cells. (A) Top view time-lapse images of apical Jup:GFP and E-Cad:3mKate2 in *mud* tissue of a reintegrating daughter cell upon a highly mis-oriented division following ablation in the apical cell center (white arrow and small dot in *t=* 0 min). Note that the apex area rapidly increased, which is considerably faster compared to the rate of reintegration without ablation (see Figure 5B). (B) Graph of the apex area change (between *t=* 0 and 79 sec) following apical cell center ablation (mean ± sem) of a reintegrating daughter cell upon a highly mis-oriented division in *mud* tissue. **** *p*<0.0001, paired student *t*-test. *n*, number of cells analyzed. (C) Top view time-lapse images of apical Jup:GFP and E-Cad:3mKate2 in *mud* tissue of a reintegrating daughter cell upon a highly mis-oriented division following apical ablation of neighboring cells (dashed white lines in *t=* 0 min). (D) Graph of the apex area change (between *t=* 0 and 79 sec) following apical ablation of the neighboring cells (mean ± sem) of a reintegrating daughter cell upon a highly mis-oriented division in *mud* tissue. **** *p*<0.0001, paired student *t*-test. *n*, number of cells analyzed. (E) Top view time-lapse images of apical E-Cad:GFP in *mud* tissue of a reintegrating daughter cell upon a highly mis-oriented division adjacent to a neighboring cell undergoing division. The apex area of the reintegrating cell (red arrowheads) increases during neighboring cytokinesis (furrow indicated by black arrows). Blue asterisks, large tissue sibling. (F) Graph of the apex area dynamics of reintegrating (small) cells adjacent to a neighboring dividing cell undergoing cytokinesis in *mud* tissue (% of the sum of the apical areas of the two siblings ± sem, sliding average 10 min). The time *t=* 0 min is set to neighboring cytokinesis onset (dashed line). *n*, number of cells analyzed. (G) Graph of the velocity of apex area expansion of reintegrating cells neighboring a dividing cell undergoing cytokinesis in *mud* tissue (µm^2^/min ± sem, sliding average 10 min). The time *t=* 0 min is set to neighboring cytokinesis onset (dashed line). *n*, number of cells analyzed. (H) Top view time-lapse images of apical E-Cad:3mKate2 of a reintegrating daughter cell (red arrowheads) upon a highly mis-oriented division in *mud zip^DN^* tissue. Note that the cell fails to reintegrate and is lost from the tissue at *t=* 220 min. Black asterisk, dividing cell at anaphase onset *t=* 0 min. Blue asterisks, large sibling. (I) Graph of the percentage of cells that initiated reintegration, but failed (as in H) in *mud w^RNAi^* and *mud zip^DN^* tissues. **** *p*<0.0001, chi-square test. *n*, number of divisions analyzed. (J) Graph of the apex area dynamics of the small daughter cells (initial apex area <10% of its sibling) in *mud* and *mud zip^DN^* tissues (mean % of the sum of the apical areas of the two siblings ± sem, sliding average 30 min). Lines indicate t_1/2_, the time to reach halfway of the predicted area (i.e. 50% of the mother apex area). The t_1/2_ of mispositioned cells is significantly different between *mud* and *mud zip^DN^* tissues. * *p*<0.05, student *t*-test. *n*, number of cells analyzed. (K) Top view time-lapse images of apical E-Cad:3mKate2 of a reintegrating daughter cell (red arrowheads) upon a highly mis-oriented division in *mud* tissue neighboring *mud* tissue over-expressing *zip^DN^* (indicated by yellow dashed lines at *t=* 0 min). Black asterisk, dividing cell at anaphase onset *t=* 0 min. Blue asterisks, large sibling. (L) Graph of the apex area dynamics of the small daughter cells (initial apex area <10% of its sibling) in *mud* tissue neighboring *mud* tissue over-expressing *w^RNAi^* or *zip^DN^* (mean % of the sum of the apical areas of the two siblings ± sem, sliding average 30 min). Lines indicate t1/2, the time to reach halfway of the predicted area (i.e. 50% of the mother apex area). The t_1/2_ of mispositioned *mud* cells abutting either *mud w^RNAi^* or *mud zip^DN^* cells is significantly different. ** *p*<0.01, student *t*-test. *n*, number of cells analyzed. (M) Graph of the apex area dynamics of the small daughter cells (initial apex area <10% of its sibling) in *mud* tissue neighboring *mud* tissue over-expressing *w^RNAi^* or *Rok^RNAi^* (mean % of the sum of the apical areas of the two siblings ± sem, sliding average 30 min). Lines indicate t_1/2_, the time to reach halfway of the predicted area (i.e. 50% of the mother apex area). The t_1/2_ of mispositioned *mud* cells abutting either *mud w^RNAi^* or *mud Rok^RNAi^* cells is significantly different. ** *p*<0.01, student *t*-test. *n*, number of cells analyzed. Scale bars: 1 µm (E, H, K), 5 µm (A, C).

Collectively, we conclude that a combination of acentrosomal MT dynamics or organization, and contractility prevent excessive cell loss upon spindle mis-orientation, and that steady-state surrounding cell contractility enhances the reintegration rate of basally mispositioned daughter cells.

### Reduced apoptosis compensates for cell loss induced by mis-oriented division

Having identified the contributions of acentrosomal MTs and contractility for cell reintegration, we next investigated the long-term consequence of cell loss due to spindle mis-orientation. Since spindle mis-orientation in *mud* scutellar tissue led to the formation of only one epithelial daughter in 27% of the divisions, it should lead to a substantial reduction in the number of epithelial cells. Given that most cells in this epithelial region of the notum undergo two rounds of division between 12 and 27 hAPF,^70,71^ we theoretically expected this cell loss to be of around 25% (see Methods). Strikingly, by comparing the tissue cell numbers at 12 and 27 hAPF in wt and *mud*, we observed that the cell number in *mud* tissue was similar to that in wt tissue (Figure 5A panel a). Importantly, in *Sas-4* tissue, spindle mis-orientation led to the formation of only one epithelial daughter cell in 31% of the divisions (*n=* 5445 divisions). Despite this excessive cell loss, the tissue cell number increase was also similar in *Sas-4* and wt scutellar tissues (Figure S5A panel a). Based on these observations, we hypothesized that a homeostatic mechanism exists to restore the tissue cell number upon cell loss due to spindle mis-orientation.

Cell loss could be compensated for by either decreased apoptosis or increased cell division. Previous studies have shown that both cell proliferation and apoptosis are spatially and temporally regulated in the notum.^70–78^ This enabled us to precisely compare cell proliferation and apoptosis rates in wt and *mud* tissues. The scutellum region is characterized by two successive waves of cell division and an increase in apoptosis rate following the first wave of cell division (Figure 5A panels b and d). During the first division wave in the tissue, the rate of cell division is similar in wt and *mud* tissue. Nevertheless, the tissue cell number increase in *mud* tissue is less than in wt tissue, in agreement with the substantial cell loss associated with mis-oriented cell division. Strikingly, the *mud* tissue cell number eventually converges to match the tissue cell number observed in wt tissue at 27 hAPF (Figure 5A panel a). Interestingly, we found that while proliferation was slightly extended in *mud* tissue after 24 hAPF, albeit at low rate, the rate of cell division producing two daughter cells was similar in *mud* and wt tissue in this extended phase (Figure 5A panel c). In contrast, we found that in *mud* tissue, the rate of apoptosis of cells within the tissue is strongly reduced following the first wave of division (Figure 5A panel d). Interestingly, quantitative analyses of cell apoptosis and division rates in *mud* and wt tissues revealed that the apoptosis rate was reduced by 4-fold in *mud* tissue and that such a reduction in apoptosis contributed to 65% of the cell number restoration (Figure S5B). As observed in *mud* tissue, we found that in *Sas-4* tissue, the tissue cell number decrease due to mis-oriented divisions is also associated with a substantial decrease in apoptosis following the first wave of division and prolonged proliferation (Figures S5A panels b and c, and S5C). Together these results prompted us to explore how the production of a lone epithelial daughter cell following a mis-oriented division could lead to a reduced rate of apoptosis to ensure that the final cell number is close to the one observed in wt tissue.

**Figure 5.**
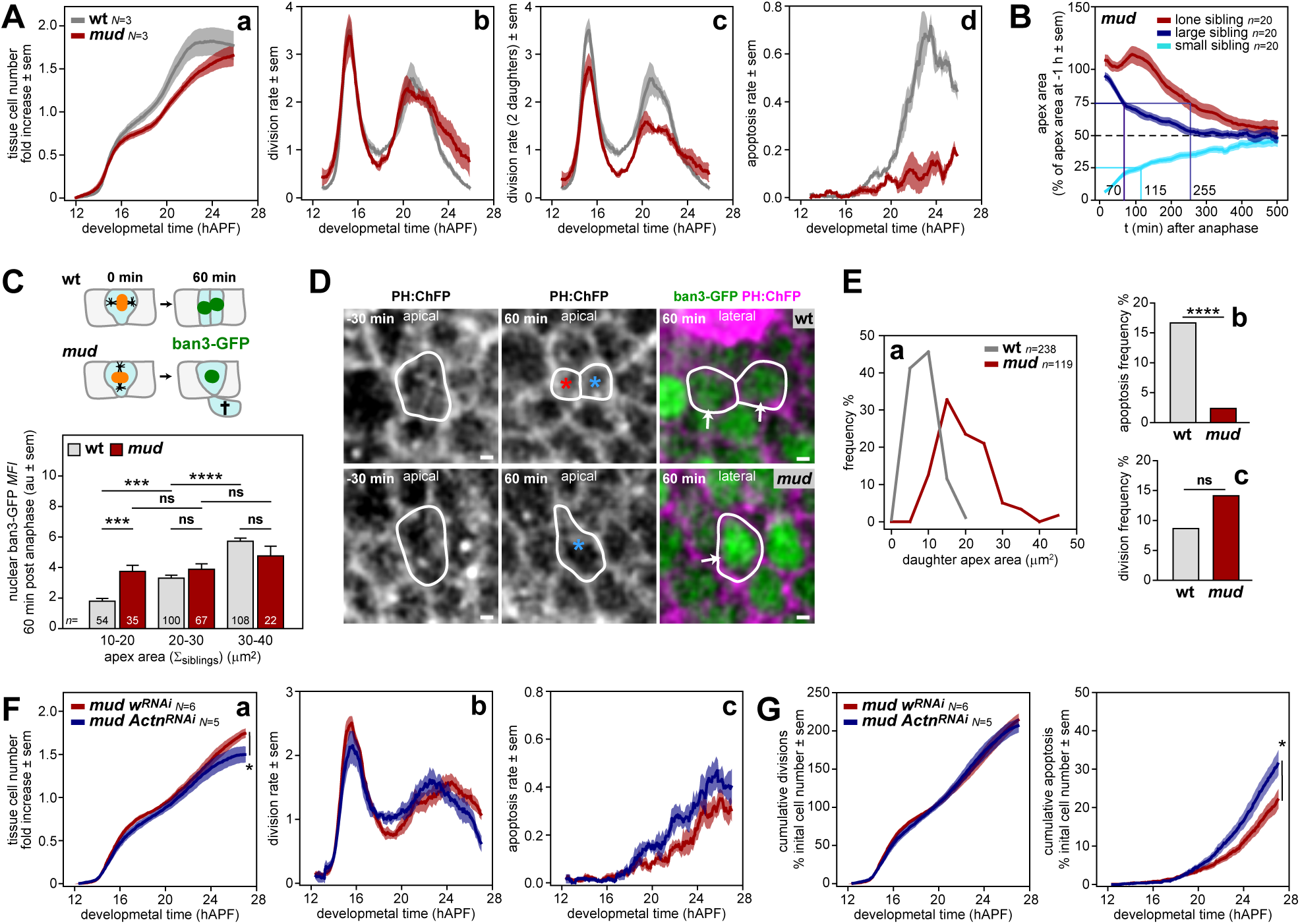
Apex area sensing by Hippo/YAP prevents apoptosis to restore tissue cell number. (A) Graphs of the tissue cell number fold increase (a)(relative to the initial cell number, mean ± sem), the total division rate (b), the division rate for divisions producing 2 daughter cells (c), and the apoptosis rate (d)(rates relative to the initial cell number, mean ± sem, sliding average 60 min) in wt and *mud* scutellum tissue. *N*, hemi-scutella analyzed. (B) Graph of the apex area dynamics of the daughter cells following *mud* divisions that produced either two epithelial daughters where one cell has and apex area <10% of its sibling (small, large) or a lone epithelial daughter. Apex areas are relative to the apex area of their mothers at -60 min prior to, *t=* 0 min anaphase onset (mean% ± sem). The time to reach halfway of the predicted area (i.e. 50% of the mother apex area) is indicated, t_1/2_ = 115, 70, and 255 min for the small, large, and lone daughters, respectively. *n*, number of divisions analyzed. (C) Schematic diagram and histogram of the quantification of the ban3-GFP nuclear mean fluorescence intensity (*MFI* ± sem) in wt daughter cells upon division, and in the lone *mud* tissue resident siblings following a highly mis-oriented division. ban3-GFP levels were measured 60 min post anaphase onset (*t=* 0 min) and cells were binned according to their initial apex area, using the sum of the siblings as a proxy (Figure S5F), to enable comparison between wt and *mud* cells. Note that in each bin the lone *mud* epithelial cells exhibit an apex area approximately twice as large as their wt counterparts. ns, not significant, *** *p*<0.001, **** *p*<0.0001, ANOVA with post-hoc Tukey test. *n*, number of divisions analyzed. (D) Top view time-lapse images of PH:ChFP at -30 min prior to and 60 min after anaphase onset (*t=* 0 min) of a dividing wt cell (top panels) and a *mud* cell producing a lone epithelial daughter (bottom panels). The images depict the apical domain of dividing cells corresponding to the smallest apical area bin (10-20 µm^2^) as quantified in C. For the 60 min time-point the lateral PH:ChFP (magenta) and the nuclear ban3-GFP (green) for the daughter cells are shown (white arrows). White outlines, cells analyzed. Blue and red asterisks indicate tissue resident daughter cells following division for which the ban3-GFP was quantified. (E) Graph of the frequency distribution (%) of daughter apex areas for wt cells after division and *mud* cells after mis-oriented divisions that produced a lone daughter cell (a), as well as the graphs of the frequencies of these wt and *mud* cells that subsequently undergo apoptosis (b) or division (c). ns, not significant, **** *p*<0.0001, chi-square test. *n*, number of daughter cells analyzed. (F) Graphs of the tissue cell number fold increase (a)(relative to the initial cell number, mean ± sem), the total division rate (b), and the apoptosis rate (c)(rates relative to the initial cell number, mean ± sem, sliding average 60 min) in *mud w^RNAi^* and *mud Actn^RNAi^* medial scutellum tissue. The final tissue cell number is significantly different between *mud w^RNAi^* and *mud Actn^RNAi^*. * *p*<0.05, student *t*-test. *N*, hemi-scutella analyzed. (G) Graphs of the cumulative divisions and apoptosis (% of initial cell number, mean ± sem) in *mud w^RNAi^* and *mud Actn^RNAi^* medial scutellum tissue. The final cumulative apoptosis is significantly different between *mud w^RNAi^* and *mud Actn^RNAi^*. * *p*<0.05, student *t*-test. *N*, hemi-scutella analyzed. Scale bars: 1 µm.

### Interplay between cell apex size and tissue mechanical tension restore cell number

To investigate the mechanisms contributing to the reduction of cell apoptosis, we focused on divisions producing only a lone epithelial daughter for two reasons. First, such divisions are the ones leading to cell loss. Second, the lone epithelial daughters were characterized by an initial apical size like the one of their mother cells for 2.67 h and subsequently decreased their apical area to half of their mother’s size (t_1/2_= 4.25 h) (Figure 5B). This contrasts with the dynamics of the daughter cells originating from highly mis-oriented divisions generating two unequal epithelial daughter cells, where the apical size reached to half their mother’s size with t_1/2_ of 1.92 h and 1.17 h for the small and large siblings, respectively (Figure 5B).

We found that the loss of daughter cells upon mis-oriented division in *mud* tissue was not associated with elevated calcium signaling (Figures S5D and S5E), indicative of the absence of a damage response associated with tissue repair upon injury.^79,80^ To identify the mechanism ensuring tissue cell number regulation, we then turned our attention to the Hippo/YAP pathway, which is a key mediator of mechanochemical signaling within tissues to control proliferation and apoptosis.^81–84^ Previously, we have shown that in the central scutellum tissue, the YAP signaling output scales as a function of apical cell size.^85^ Furthermore, the rate of apoptosis inversely correlates with cell apical area and YAP activity.^72^ In wt tissue, each division produces two daughter cells whose apical areas are close to half the size of their mother cell.^71,72^ In contrast, in *mud* tissue, with mis-oriented cell division producing only one epithelial daughter, the apical area of this lone epithelial sibling is similar to that of its mother (Figure S5F). Therefore, cell loss in *mud* tissue is concomitant with the formation of a lone sibling having an apex area twice as large as that observed in wt tissue. Given that YAP activity scales with apical area in the regulation of cell apoptosis,^72,85^ we hypothesized that the production of only one epithelial daughter cell exhibiting a larger apical size could account for the reduced apoptosis in *mud* tissue.

Hippo/YAP signaling modulates apoptosis through the transcriptional control of the microRNA bantam (ban),^86,87^ whose expression can be used as a readout for pathway activity using the ban3-GFP transgene.^88^ To test whether Hippo/YAP signaling could participate in the change in apoptosis in *mud* epithelial tissue, we measured the ban3-GFP level in wt and *mud* cells. Specifically, one hour after division, we compared the nuclear ban3-GFP level in wt daughter cells and in *mud* daughter cells originating from divisions that produced only a lone epithelial daughter cell. Importantly, to compare the ban3-GFP level in wt and *mud* cells, cells were binned according to their mother’s apical cell size, using the sum of the apical areas of the siblings as a proxy (Figure S5F). As observed when measuring YAP activity in wt tissue at any time point during the cell cycle,^85^ we found that the ban3-GFP intensity scales with apex area in wt cells as early as 1 h after cell division (Figures 5C and 5D). In wt tissue, the offspring from the 10-20 µm² mothers constitute the smallest cells in the tissue; they have the lowest YAP activity and the highest probability of undergoing apoptosis.^72^ Strikingly, the ban3-GFP intensity in lone *mud* epithelial daughters of the smallest mother cells was twice as high as in wt offspring originating from the smallest mother cells (Figure 5C). Therefore, in *mud* tissue, the lone epithelial daughter cells originating from the smallest sized mothers harbor an apical area twice as large as their wt counterparts and are characterized by a higher level of Hippo/YAP signaling.

To investigate whether this increase in YAP activity correlated with a lower rate of apoptosis, we tracked the fate of the lone *mud* daughter epithelial cells alongside wt daughter cells of comparable mother apex size. Strikingly, we found that the frequency of *mud* cells originating from divisions producing only one epithelial daughter cell to undergo subsequent apoptosis was drastically reduced compared to that observed in wt cells (Figure 5E panels a and b). Furthermore, in line with the slight change in the rate of cell division in *mud* tissue, the frequency of such cells undergoing division was elevated, although not significantly compared to wt cells (Figure 5E panel c). Therefore, we conclude that the reduced rate of apoptosis in *mud* tissue is associated with the formation of daughter cells that are similar in size to their mother and exhibit a higher level of Hippo/YAP signaling as well as a lower rate of apoptosis.

Since the scaling between Hippo/YAP activity and apical cell size depends on the actin cross-linker α-Actinin (Actn)^85^, we reduced Actn function in *mud* (*mud Actn^RNAi^*) and quantified the tissue cell dynamics. In *mud Actn^RNAi^*, the proliferation rate was largely unaffected, but apoptosis was significantly increased, resulting in a decrease in tissue cell number compared to *mud w^RNAi^* (Figures 5F, 5G, S5G and S5H), which was not due to a change in the Rd/r ratio (Figure S5I). This is consistent with a model in which sensing of apex area in response to tissue tension by Hippo/YAP pathway prevents apoptosis in cells with relatively larger apices. Altogether, we conclude that upon spindle mis-orientation the decrease in cell apoptosis within the epithelial tissue is contributing to cell number homeostasis. Furthermore, we propose that this decreased apoptotic rate is due to the formation of only one lone epithelial daughter harboring an apical size equal to that of its mother, leading to higher YAP signaling.

### Systemic TNF signaling eliminates basally displaced cells that fail to reintegrate

To further challenge this model, we wondered whether the death of daughter cells positioned below the tissue could contribute to regulating epithelial cell numbers. To address this question, we decided to identify the mechanisms that promote their death to prevent it. We hypothesized that upon mis-oriented division the basally located cells lose their epithelial polarity and epithelial barrier function. Accordingly, basally born cells lack a polarized distribution of the apical marker GFP:aPKC (Figure S6A). Therefore, we focused on the role of systemic Tumor Necrosis Factor (TNF) signaling, which has recently been shown to control the elimination of polarity-deficient epithelial cells.^89^ Since, the *Drosophila* TNF ligand, Egr,^90,91^ is produced by the fat body cells and secreted into the hemolymph,^92^ we first downregulated Egr specifically in the fat body cells using the *lpp-GAL4*^93^ driver in wt and *mud* animals. Quantification of cleaved Casp3 positive cells beneath the tissue showed that while *egr^RNAi^* in the fat body did not change the number of basal apoptotic corpses in control tissues, *egr^RNAi^* in the fat body of *mud* animals strongly reduced the apoptosis observed beneath *mud* epithelial tissue (Figures 6A and 6B). Secondly, we downregulated the *Drosophila* TNF receptor Grnd^94^ in wt and *mud* tissues to test whether this would prevent apoptosis of the basally born cells. Indeed, like the depletion of *egr^RNAi^* in the fat body, compared to *mud w^RNAi^* control tissue the number of basal apoptotic corpses was strongly reduced in *mud grnd^RNAi^* (Figures 6C and 6D). Furthermore, tracking of the basally born cells, labelled using the DNA replication marker PCNA:GFP,^95–97^ revealed that knockdown of *grnd* in *mud* tissue led to 26% (5/19) of the basally located cells to re-enter the cell cycle, as they underwent S-phase to then likely remain in G2 (Figures S6B-D). This also confirmed that *grnd^RNAi^* in *mud* tissue is necessary to promote the apoptosis of basally located cells since 30% of the basally delaminated cell did not exhibit signs of apoptosis as compared to all 17 tracked control *mud w^RNAi^* cells that showed nucleus fragmentation (Figure S6C). Importantly, *grnd^RNAi^* in *mud* tissue neither modulated the R_d/r_ ratio, nor changed the proliferation and apoptosis rates, nor modified the tissue cell number (Figures S6E and S6F). We therefore uncovered that the elimination of basally born daughter cells upon mis-oriented division is promoted by systemic TNF signaling and that their elimination is not necessary to regulate epithelial cell number. Altogether, our work establishes how complementary cell, tissue, and systemic mechanisms act to control cell number within and outside the epithelium upon spindle mis-orientation.

**Figure 6.**
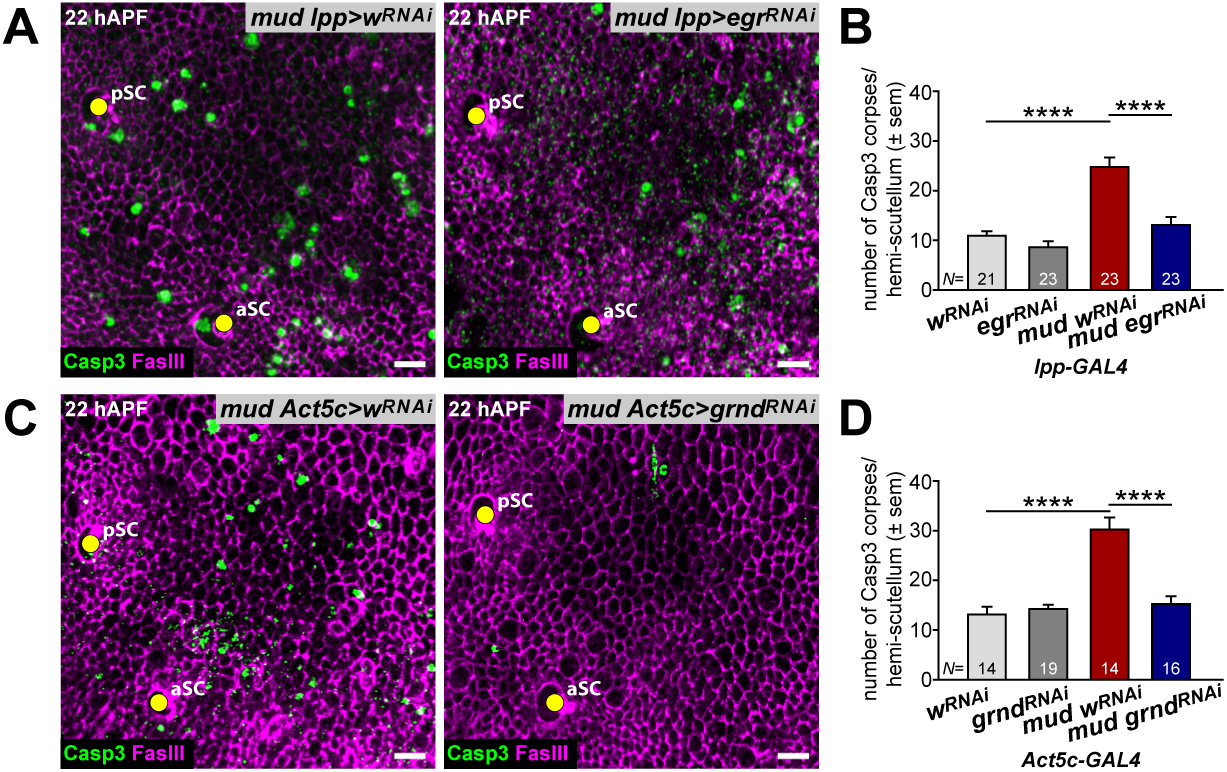
Systemic TNF signaling eliminates misplaced basal cells. (A) Top view images of fixed *mud lpp>w^RNAi^* and *mud lpp>egr^RNAi^* tissues at 22 hAPF stained with antibodies against cleaved Casp3 and against SJ marker FasIII. aSC and pSC, anterior and posterior scutullar macrochaetes. The *lpp-Gal4* driver specifically expressed the RNAi in the fat body. (B) Graph of the number (mean ± sem) of Casp3 positive corpses per hemi-scutellum tissue in wt and *mud* tissues of *lpp>w^RNAi^* and *lpp>egr^RNAi^* animals. **** *p*<0.0001, ANOVA with post-hoc Tukey test. *N*, number of tissues analyzed. (C) Top view images of fixed *mud Act5c>w^RNAi^* and *mud Act5c>grnd^RNAi^* tissues at 22 hAPF stained with antibodies against cleaved Casp3 and against SJ marker FasIII. aSC and pSC, anterior and posterior scutullar macrochaetes (yellow circles). The *Act5c-Gal4* driver promotes ubiquitous expression of the RNAi. (D) Graph of the number (mean ± sem) of Casp3 positive corpses per hemi-scutellum tissue in wt and *mud* tissues of *Act5c>w^RNAi^* and *Act5c>grnd^RNAi^* animals. **** *p*<0.0001, ANOVA with post-hoc Tukey test. *N*, number of tissues analyzed. Scale bars: 10 µm.

## Discussion

Tissue cell number and positioning are tightly regulated during epithelial development, homeostasis, and repair. Here, we induced cell mispositioning and loss by disrupting spindle orientation to investigate the mechanisms that maintain tissue architecture and cell number during development. We established the role of the MT and actomyosin cytoskeletons in regulating the balance between cell reintegration and delamination in mispositioned cells. Additionally, we elucidated that apical cell size and Hippo/YAP signaling promote the restoration of cell number within the tissue upon cell loss. Lastly, we found that systemic TNF signaling selectively promotes the death of delaminated cells, hindering the accumulation of potentially harmful cells. Thus, we propose that epithelial integrity and cell number upon spindle mis-orientation rely on complementary functions of local tissue mechanics and systemic signaling, collectively safeguarding the tissue and the organism against mispositioned cells and cell delamination. Our work positions the investigation of spindle mis-orientation as a paradigm for exploring how cell-, tissue-, and organism-scale processes maintain tissue architecture and cell number.

Cell reintegration is crucial for preventing excessive cell loss upon spindle mis-orientation and has also been identified as a physiological process associated with cell dissemination.^8,98,99^ The mechanisms regulating cell reintegration upon division mis-orientation have predominantly been studied in *Drosophila* follicular cells, where all cells reintegrate in an apical-to-basal fashion. In this context, lateral SJ cell-cell adhesion proteins promote reintegration.^13,14^ In other tissues and in the notum, reintegration occurs from basal-to-apical.^12,24^ Whether these different modes of cell reintegration rely on similar mechanisms has been unclear. We found that lateral SJ, TCJ, and E-Cadherin adhesion molecules are not required for basal-to-apical cell reintegration, and identified the cytoskeleton’s contributions in both the reintegrating cell and its neighbors. Firstly, we established that acentrosomal Pat- and Eb1-dependent MTs are necessary for maintaining the apical cap associated with the reintegration of basally mispositioned cells. Given that MT spatial organization and dynamics regulate actomyosin contractility,^100–102^ membrane protein delivery,^48,103,104^ and pushing forces on the membranes,^105–107^ these functions likely contribute to the stabilization or maintenance of the apical cap during the initial step of cell reintegration. Additionally, in line with previous studies demonstrating the importance of Pat distribution on MT network organization,^42,46,49,51–53,108^ we found that loss of the apical transmembrane protein Sas led to both the delocalization of Pat from the apical medial cortex to the cell junctions and altered the reintegration of mispositioned cells. This suggests that the organization of the apical acentrosomal cytoskeletal MT network is tuned to ensure cell reintegration. Exactly, how Sas regulates Pat localization remains to be further investigated, but might involve its receptor PTP10D, whose trans-interaction with neighboring cells has been shown to eliminate potentially tumorigenic cells.^58–60,109^ Secondly, we found that surrounding epithelial cells promote reintegration of mispositioned cells by exerting pulling forces. These forces, which originate from the steady-state tensile forces within the epithelium, are necessary to prevent delamination and enhance the rate of cell reintegration. Our work thus suggests a mechanical interplay between mispositioned cells and their surroundings during reintegration. While acentrosomal Pat- and Eb1-dependent MT function in apical cap maintenance is restricted to the early step of the cell reintegration process, contractility also supports tissue maintenance and regulates cell shape, as well as their variation, throughout tissue development.^110,111^ Previous work on *Xenopus* radial cell intercalation has shown the critical roles of TCJ, actomyosin forces, and centrosomal MTs in promoting the emergence of basally located cells.^27,28,112^ Our analysis revealed that, despite similarities in cell dynamics, the regulation of basal-to-apical cell reintegration of basally mispositioned cells differs from radial intercalation. Specifically, while radially intercalating cells push their neighbors, we found that mispositioned cells are rather pulled by surrounding cells. Furthermore, centrioles play a crucial role in radial cell intercalation,^38^ yet centrosomes were not essential for reintegration of mispositioned daughter cells in the notum epithelium. Last, our study also highlights the existence of mechanisms controlling the relative apical size of daughter cells within a tissue. Upon spindle mis-orientation, daughter cells with unequal apical sizes rapidly adjust their apex areas to match those observed under normal conditions. The mechanisms controlling apical cell area and its relation to cell survival remain to be understood, and spindle mis-orientation could be used to further probe the contributions of contractility, cell-cell adhesion, cell-extracellular matrix adhesion, apical-basal polarity, and cell-cell signaling.

During development, homeostasis, and repair, the number of cells within tissues is tightly controlled by the balance between cell proliferation and death. Classical approaches to probing the mechanisms regulating epithelial cell number have relied on inducing local changes in growth, proliferation, or apoptosis rates. This has enabled the identification of key mechanisms that safeguard tissue cell numbers, such as cell competition and compensatory cell proliferation upon cell apoptosis.^113–123^ In addition, mechanical stress, which can propagate over long distances, has emerged as a potential long-range regulator of cell proliferation and apoptosis in epithelial cell density control.^124–128^ However, it remains critical to understand how these diverse processes quantitatively account for the global control of tissue cell numbers upon cell loss. Here, we established that the loss of one daughter cell upon spindle mis-orientation leads to the formation of a lone epithelial daughter with an apical size similar to that of its mother. Furthermore, these lone daughter cells exhibit higher levels of YAP activity and a reduced rate of apoptosis. This regulation predominantly affects the smallest lone daughter cells, consistent with our previous findings that Hippo/YAP signaling is controlled by apical area up to a certain apical area threshold.^72,85^ Since we previously found that Hippo/YAP signaling is dependent on global tissue mechanical stress and cell apical size, our findings delineate how the complementary roles of tissue mechanical stress and cell geometry can locally adjust cell number. This regulation might support the hypothesis that cell apical area, rather than cell volume, is a factor in controlling the number of cells within a tissue. Importantly, our data also suggest that the observed increased rate of cell division upon loss of Mud function could result from the additional proliferation of lone daughter cells. Interestingly, the major change in ban3-GFP level between wt daughters and *mud* lone epithelial daughters occurs for the *mud* lone daughter cells of apex area lower than 20 µm^2^. This is rather expected since we previously found,^85^ that the scaling between apical size and ban3-GFP mainly happens for apex area between 5 and 25 µm^2^. The loss of Mud function therefore leads to changes in ban3-GFP level and apoptosis in the smallest cells of the tissue, which are the ones prone to die in wt tissue.^72,85^ As compensatory cell proliferation has been proposed to safeguard against spindle mis-orientation in other tissues,^7^ it will be relevant to explore the roles of Hippo/YAP signaling, cell apical area, and mechanical stress in response to spindle mis-orientation more broadly. More generally, the complementary inputs provided by global tissue mechanics and cell geometry are emerging as a paradigm for locally modulating cell number within tissues. In collective cell assemblies *in vitro*, it has been found that both mechanical forces and cell area are good predictors of cell proliferation,^129^ and that local tissue tension and nucleus size induce the proliferation of a subset of cells surrounding apoptotic cells.^123^ It will be interesting to further explore how nucleus size and cell area are correlated in tissues and how their respective contributions are integrated to control proliferation and apoptosis, thereby robustly controlling tissue cell number.

While Hippo/YAP activity emerges as a regulator to prevent the death of large lone daughter cells upon spindle mis-orientation, we foresee that ERK and Notch signaling, or Hippo/YAP-mediated compensatory proliferation could act in parallel to the anti-apoptotic function of the Hippo/YAP pathway. First, we previously uncovered that Notch signaling is critical for modulating apoptosis as a function of relative cell apical area.^72^ It is therefore possible that the larger cell size of lone daughter cells relative to their direct neighbors also prevents their death. Second, the ERK pathway has recently emerged as a regulator of cell apoptosis in response to local mechanical forces, dependent on the pulling forces associated with the delamination of dying cells.^73,130^ Since, upon spindle mis-orientation, the lone daughter cell initially occupies the space of the mother, spindle mis-orientation is unlikely to initially promote ERK activation. Nevertheless, over longer time scales, the large lone daughter decreases its area, possibly pulling on its neighbors and increasing ERK activity to prevent the death of its surrounding cells. Finally, the mechanism we uncovered can only compensate for epithelial cell loss if the endogenous apoptosis rate is higher than the rate of cell loss induced by spindle mis-orientation. If the endogenous rate of apoptosis is too low, our data suggest that the rate of proliferation could be enhanced to ensure cell number regulation, possibly via Hippo/YAP signaling. This could explain why higher levels of cell proliferation were reported in *Drosophila* epithelial tissue lacking centrosomes, which induced cell loss via spindle mis-orientation, DNA damage and aneuploidy.^7,131^ As cell proliferation requires cell cycle progression, it is possible that a reduced rate of apoptosis acts as an early response, while compensatory proliferation can act on a longer time scale in the case of more significant cell loss.

While spindle mis-orientation can predispose basally born epithelial daughter cells to initiate tumor formation upon apoptosis inhibition,^8^ the mechanisms selectively inducing the death of delaminated cells have remained unclear. We propose that the loss of cell polarity in delaminated cells signals their elimination. Loss of cell polarization, caused by the loss of key polarity factors or tumor suppressors, is known to activate the TNF pathway due to mis-localized TNF receptors activated by the systemic TNF ligand Egr.^89^ We showed that the TNF ligand Egr, produced by the fat body, promotes the death of delaminated cells. This function is limited to mispositioned cells, suggesting that loss of polarization is crucial for selective cell death. Furthermore, the death of delaminated cells does not affect epithelial apoptosis or proliferation rates, indicating that delaminated cell death is unlikely to signal for regulating epithelial cell numbers. Apoptosis prevention has been shown to induce cellular senescence and to arrest cells either at the G1 or G2 phase of the cell cycle.^132–135^ Similarly, we found that loss of the TNF receptor Grnd reduced basal cell apoptosis, and that some basally born cells were able to progress through S-phase. Because we did not observe any basal cell division, these data indicate that these cells might remain in G2 phase. Since in numerous contexts TNF regulates JNK signaling, which mediates stress-induced apoptosis,^89,136–140^ we envision that Egr promotes the death of basally located daughter cells via activation of the JNK pathway. Previous work showed that loss of centrosomes causes cell apoptosis in a JNK-dependent manner, but the downregulation of *Egr* expression in this epithelium did not prevent apoptosis.^7^ Building on our findings, it will be interesting to test the role of systemic Egr in controlling cell death induced by centrosome loss. Collectively, our results indicate that the systemic TNF pathway enables for detecting delaminated cells in the absence of gene function loss that regulate epithelial polarity, delineating a simple and potentially common mechanism preventing the accumulation of non-epithelial, cancer-prone cells.

Development entails the highly complex spatiotemporal dynamics of thousands of cells. It has emerged as an error-prone process backed-up by compensatory and checkpoint mechanisms. Loss-of-function analyses have been instrumental in delineating such mechanisms that ensure the robustness of tissue development and have provided key insights into tissue homeostasis, repair, and aging. This includes: at the cellular level, mitotic checkpoints ensure the fidelity of genome segregation^141^; at the tissue level, cell competition and compensatory proliferation buffer local changes in growth, apoptotic rates, or cell identities^113–122^; or at the organismal level, inter-organ communication helps maintain organ allometry and symmetry.^142,143^ Spindle mis-orientation is associated with numerous pathologies and arises from defects in cortical cues, cell contractility, and microtubule organizing centers.^4,8,144^ Our study reveals a multilayered system of safeguards that reuse robust homeostatic mechanisms to maintain epithelial integrity and cell number in the face of spindle mis-orientation, opening new avenues for investigating the complex interplay between cell behavior, tissue mechanics, and systemic regulation in epithelial biology.

### Limitations of the study

The microtubule-associated proteins (MAPs) Pat and Eb1 have important functions in the regulation of MT dynamics and organization. Our findings regarding the function of these MAPs suggest that their activity is confined to mispositioned reintegrating cells to prevent their loss. However, we cannot formally rule out a function in neighboring cells during cell reintegration. In addition, the limits imposed by live imaging impeded us from delineating whether they act by controlling MT dynamics or organization in the apical cap of the reintegrating daughter cells. To investigate the role of Hippo/YAP signaling we analyzed the ban3-GFP at one hour post division. This analysis required the simultaneous acquisition of the 3D cell outlines using PH:ChFP with ban3-GFP. Photobleaching and 3D cell tracking hindered the quantitative analyses of the ban3-GFP level changes over time to explore the dynamics of Hippo/YAP signaling in lone daughter cells upon spindle mis-orientation. Due to similar challenges in live cell imaging and tracking, cell cycle progression of basally born cells using the PCNA:GFP marker combined with PH:ChFP could only be reliably followed for an average of 4.2 ± 0.2 h after a mis-oriented division. Although all basally born cells underwent apoptosis in control *mud w^RNAi^* tissue, due to the technical limitation to track for extensive periods the basally born cells in *mud grnd^RNAi^* cells, further work will be necessary to fully characterize their fate and tumorigenic potential.

## STAR METHODS

## RESOURCE AVAILIBILITY

### Lead contact

Further information and requests for resources and reagents should be directed and will be fulfilled by the lead contacts, Floris Bosveld and Yohanns Bellaïche.

### Materials availability

All unique/stable regents generated in this study are available from the lead contacts without restriction.

### Data and code availability

- All original microscopy data reported in this manuscript will be shared by the lead contacts upon request.
- Any additional information required to reanalyze the data reported in this manuscript is available from the lead contacts upon request.
- All code associated with this study will be shared by the lead contacts upon request.

## EXPERIMENTAL MODEL AND SUBJECT DETAILS

### *Drosophila melanogaster* husbandry and stocks

Fly stocks were maintained on standard molasses/cornmeal/yeast food at 18°C or 25°C. Experiments were performed at 25°C or 29°C as specified in the Methods. The *Drosophila* stocks used in this study are listed in the Key resource table.

## METHODS DETAIL

### Genetics

Experiments were performed at 25°C, or 29°C when utilizing the *Act5c-GAL4* driver for global over-expression experiments. For temperature-controlled experiments we used the GAL4/GAL80^ts^ system^145^ and these animals were reared at 18°C. To induce expression animals were pre-incubated at 29°C for 24-30 h prior to imaging, which generally commenced at 12-15 hours after pupa formation (hAPF). For clonal analyses, the flip-out system was utilized^146^. Thereto larvae were exposed to a 37°C heat-shock (7-20 min) and imaged 3-4 days after heat-shock.

### Construction of Ck:GFP

The Ck:GFP allele was generated by CRISPR/Cas9-mediated homologous recombination at its endogenous locus, using the vas-Cas9 line.^147^ Thereto guide RNAs: 5’-GTCGTATGTGTAGGTTTTACTAGTGTTT-3’ and 5’-GTCGAAATATTTTGCGTTATGTGTGTTT-3’, were cloned into the pCFD5:U6:3-t::gRNA vector.^148^ The homology sequences: HR1 and HR2 were amplified using the following primers: HR1F-5’-TATGGGGTGTCGCCCTTCGGGTCTCTAGTGTAAAGATCGCGGATAAGGTCA-3’, HR1R-5’-ACTGCCTGAAGAACCGCTGGACCCCGAACTGTTGGCTCGAATGGTTCGATT-3’; HR2F-5’– GGAAGTGGTAGCTCAGGGTCTAGTGGATAGCACATAACGCAAAATATTT-3’, HR2R-5’-GCCCTTGAACTCGATTGACGCTCTTCGTCCGTCAGGCAAGAAAATCAAGGGA-3’ and cloned, using SLIC,^149^ into a recombination vector, p931_GFPCter_Mini-White, carrying a hs-mini-white cassette flanked by two LoxP recombination sites for its removal and a C-terminal GFP.^150^ All constructs were confirmed by sequencing and embryo transgenesis was performed by Bestgene.

### Immunostaining of fixed tissue

Pupae were prepared for fixed tissue imaging as described previously.^151^ Tissues were stained with primary antibodies: rabbit anti-cleaved Caspase3 (1:200), guinea-pig anti-Cora (1:2000), mouse anti-FasIII (1:50), rat-anti-αCat (1:250), mouse anti-BrdU (1:25) and subsequently counterstained with fluorescently tagged secondary antibodies (1:200): goat-Alexa488 anti-rabbit, donkey-Cy5 anti-guinea-pig, donkey-Cy3 anti-mouse, goat-Alexa555 anti-rat.

### BrdU incorporation assay

BrdU staining was performed by dissecting notae (14-16 hAPF) from PCNA:GFP expressing animals in Schneider’s *Drosophila* medium. Notae were subsequently cultured in Schneider’s *Drosophila* medium containing 75 µg/ml BrdU for 30 min at 25 °C, washed two times 5 min with Schneider’s *Drosophila* medium and once 5 min with 1xPBS, after which the tissues were fixed for 20 min in 1xPBS containing 4% PFA. After two 10 min washes in 1xPBS + 0.3% Triton X-100, and two 10 min washes in 1xPBS + 0.5% Triton X-100, notae were permeabilized for 40 min in 1xPBS + 0.5% Triton X-100. Next, the DNA was depurinated for 30 min in a 1:1 mixture of 1xPBS + 0.5% Triton X-100: 4N HCl (2N final concentration), after which tissues were washed three times 10 min in 1xPBS + 0.25% Triton X-100. Immunolabelling and mounting was performed as described in immunostaining of fixed tissue.

### Image acquisition, processing, and quantification

Mounting of pupae for live imaging was performed as in described in.^71^ Microscopes used in this study for fixed or live tissue imaging include: an inverted spinning disk wide borealis confocal microscope CSU-W1 (Andor/Roper/Nikon) with sCMOS camera (Orca Flash4, Hamamatsu), an inverted spinning disk wide homogenizer confocal microscope CSU-W1 (Roper/Zeiss) with sCMOS camera (Orca Flash4, Hamamatsu), an inverted spinning disk wide confocal microscope CSU-W1 (Roper/Nikon/GATACA) with sCMOS camera (BSI camera with 95%QE), an inverted spinning disk wide homogenizer confocal microscope CSU-W1 (Roper/Zeiss/Spark) with sCMOS camera (Orca Flash4, Hamamatsu), an inverted spinning disk confocal microscope (Roper/Nikon) with sCMOS camera (Orca Flash4, Hamamatsu), and a spinning disk upright confocal microscope (Roper/Zeiss) with Coolsnap HQ2 camera (Photometrics).

#### Laser ablations

For tissue laser ablations, we used an inverted laser scanning microscope (LSM880 NLO, Carl Zeiss) equipped with a multi-photon Ti::Sapphire laser (Mai Tai HP DeepSee, Spectra-Physics). Circular and point ablations were performed in tissue expressing the microtubule associated protein Jup:GFP as well as the E-Cad:3mKate2 AJ marker by using a 40x/1.3 OIL DICII PL APO (UV) VIS-IR (420762-9800) objective and imaging in single-photon bidirectional scan mode with 1.0 sec time-steps. To evaluate the apex area change upon ablation, a small point was ablated in the medial cortex of the reintegrating cell, or a region of interest (ROI) drawn around the reintegrating cells was severed using the Ti::Sapphire laser in two-photon mode at 820-890 nm. The change in apex area was measured between *t_0_=* 0 and *t_80_=* 79 sec and statistical significance evaluated using the paired *t*-test. For junction ablations we used a 63x/1.4 OIL DICII PL APO objective and imaged a region of 26.45 µm x 26.45 µm (301x301 pixels). Junction ablations were performed in tissue expression the AJ marker E-Cad:GFP and imaged with 0.56 sec time-steps. To estimate the relative junctional tension,^152^ we measured the recoil velocity of the vertices after junction ablation between *t_0_*= 0 sec and *t_7_*= 3.92 sec. Ablations were performed in *mud* tissue to probe the relative tension in the junctions of cells neighboring a reintegrating cell (+1N) as well as the relative local tissue tension by ablation of junctions belonging to cells 2 cell distances away from the reintegrating cell (+2N)(Figure S4C). To enable comparison, we ablated in a single experiment two junctions, alternating between either first a +1N junction or a +2N junction. For each experiment we determined the ratio of the measured relaxation velocities, +1N/+2N. Statistical significance was evaluated using the paired *t*-test. Similar ablation experiments were performed to estimate the relative junction tension in +1N cells in *mud w^RNAi^* and *mud pat^RNAi^* double mutant tissues.

#### Delamination versus reintegration

In all analyses delamination and reintegration events were visually identified and quantified in Fiji^153^ on Z-stack live-images of tissues expressing either AJ markers ubi-E-Cad:GFP, E-Cad:GFP, E-Cad:3mKate2 or the SJ marker Dlg:GFP. Typically 16-26 Z-stack slices at 0.5 µm spacing per time-point were acquired with a temporal resolution of 2-5 min between 12 to 28 hAPF using ojectives: 63x/1.4 OIL DICII PL APO, 60x/1.4 OIL DIC N2 PL APO VC, 40x/1.4 OIL DIC H/N2 PL FLUOR, 40x/1.3 OIL DIC H/N2 PL FLUOR, 40x/1.3 OIL DIC PL APO (UV) VIS-IR. Both events were scored following mitotic rounding/apex area expansion. In all analyses the anaphase elongation and apex area reduction after maximal rounding and expansion was set to reference *t=* 0 min. All data plotted and analyzed for reintegrating daughter cells represents daughter cells exhibiting an apex area <10% of the sum of the two siblings. Delaminations were scored by rounding/apex area expansion following the production of a lone epithelial sibling.

Scoring the delaminating versus reintegrating cells within the tissue yielded the ratio R_d/r_, which was used to evaluate differences amongst genotypes, and to evaluate whether dsRNA mediated knock-down in mosaic tissues generated in a *mud* mutant background modulates cell reintegration by comparing the R_d/r_ in *mud* cells and the neighboring clonal double mutant tissue. For these analyses similar numbers of divisions were analyzed between 12 to 18 hAPF in the *mud* and double mutant tissue. The data from multiple animals was pooled and the R_d/r_ in *mud* cells was set to 1 for each condition analyzed. To compare the proportions across the different categories (e.g. *mud* and surrounding double mutant tissue), we used the chi-square test to evaluate statistical significance. Similar clonal analyses were performed in wt tissue; at least 100 divisions were analyzed for each condition. None of the dsRNA transgenes reported in this manuscript elicited a phenotype in wt tissue, indicating their specificity in modulating epithelial cell delamination and reintegration only in context of mispositioned cells upon division mis-orientation.

#### Dynamics of lone epithelial daughters and reintegrating cells

To follow the apex area dynamics of the two daughter siblings upon a highly asymmetric division and of the lone epithelial daughter after division mis-orientation, their apex areas were measured in Fiji. The resulting areas were normalized to the apex area of their mother at -1 h prior to division and plotted as a 30 min sliding average. Similar quantifications were performed to compare the reintegration dynamics of mispositioned daughter cells in different genotypes, by plotting the apex area of the small reintegrating sibling as a percentage normalized by the sum of the two siblings. The time to reach halfway of the predicted area (i.e. 50% of the mother apex area), denoted as t_1/2_, was used to evaluate whether reintegration dynamics differed between genotypes, using a student *t*-test.

#### Mitotic spindle orientation

Spindle orientation along the apical-basal (αAB) tissue plane was determined for dividing cells between 12 to 18 hAPF, at *t=* -2 min prior to anaphase elongation, using Fiji and a custom plugin, in tissue expressing either a centrosome (YFP:Asl, Spd-2:RFP) or mitotic spindle (αTub:GFP, RFP:αTub) marker, as previously described.^20^ Z-stack live-images (40-50 slices at 0.5 µm spacing) were aquired with a temporal resolution of 2 min using either a 100x/1.45 OIL PL APO L, a 100x/1.4 OIL DIC N2 PL APO VC, a 63x/1.4 OIL DICII PL APO or 60x/1.4 OIL DIC N2 PL APO VC objective.

To evaluate how AB spindle orientation related to the production of either a lone epithelial daughter or resulted in a division producing a daughter cell with a small apex that reintegrated back within the epithelial tissue plane, we imaged *mud* tissue expressing the E-Cad:GFP AJ, the centrosomal YFP:Asl and RFP:αTub mitotic spindle markers using a 63x/1.4 OIL DICII PL APO objective and aquired Z-stack images (40 slices at 0.5 µm spacing) with a temporal resolution of 2 min. Statistical significane was evaluated using the Mann-Whitney U test.

#### Tissue cell number, division, and apoptosis rates

To quantify tissue cell number increase, and the division and apoptosis rates, Z-stack live-images (26 slices at 0.5 µm spacing) of tissues expressing AJ markers ubi-E-Cad:GFP, E-Cad:GFP or E-Cad:3mKate2 were acquired with a temporal resolution of 5 min between 12 to 28 hAPF using a 40x/1.4 OIL DIC H/N2 PL FLUOR, a 40x/1.3 OIL DIC H/N2 PL FLUOR or a 40x/1.3 OIL DIC PL APO (UV) VIS-IR objective. To keep the focus of the tissue, autofocus detection (Metamorph software) was used for the duration of the imaging. The aquired movies were processed and projected (maximum intensity Ζ-stack projection) using a CARE deep neural network model^154^ adapted to the notum tissue.^72^ All movies were temporally aligned using ¾ of the first division peak.^71^ The ROI for quantification, which corresponds to the medial scutellum tissue and region previously analyzed,^85^ was defined by a squared box whose exterior was delineated by the positions of the posterior dorsocentral (pDC), the anterior and posterior scutullar macrochaetes (aSC and pSC), as well as the axis of symmetry, the tissue midline. Within this ROI cells were manually tracked to quantify divisions, cell reintegration or cell delamination caused by mis-oriented cell division or apoptosis using Fiji. To quantify tissue cell number, and the division and apoptosis rates within the entire scutellum tissue the ROI was defined by the midline and the pDC, pSC and the anterior postalar (aPA) macrocheates.

For all analyses of cell number changes, division and apoptosis rates, the scored number of events were normalized to the initial number of cells tracked within the tissue. To evaluate statistical significance, we performed a student *t*-test on the final time point when comparing their cumulative effects.

To evaluate the fate of lone epithelial daughter following mis-oriented divisions, we tracked these cells, which were born during the second division wave, during a period of 12.16 h starting at 19 hAPF. We recorded their subsequent division or apoptosis. For comparison with wt daughter cells following division, we selected an equal number of cells with similar apex area from an equal number of tissues, ensuring their positions corresponded to identical locations to those analyzed in *mud* tissue within the analyzed ROI. The chi-square test was used to evaluate statistical significance.

#### Estimation of the contribution of the changes in cell division orientation as well as cell division and apoptosis rates in the variation of cell number in control and mud tissues

Within a tissue where all cells divide twice and 27% of cell divisions in *mud* produced only one epithelial daughter, the cell number is expected to be reduced by 25% relative to the control tissue. Indeed, if we consider a tissue initially composed of n cells, 𝑛*_cells_* in wt tissue, the cell number upon one division is 2𝑛*_cells_*, and the number of cells upon two divisions is 4n cells. In *mud* tissue, the cell number upon one division is 𝑛*_cells_* + 𝑛*_cells_*\* 0.73 = 1.73𝑛*_cells_*. Upon the second division, the number of cells is 1.73𝑛*_cells_* + 1.73𝑛*_cells_* * 0.73 ≈ 3𝑛*_cells_*. Therefore, the cell number in *mud* tissue is 75% of the number of cells in wt tissue.

To account for the experimental division rate, the apoptotic rates as well as to estimate the final number of cells in *mud* tissue if no compensatory process occurred, we turned to computational analyses of the contribution of division and apoptosis to the variation of cell number fold change.

We first considered that the tissue cell number evolves as:

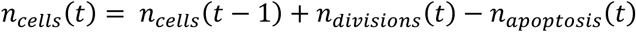

Thus, by mathematical induction:

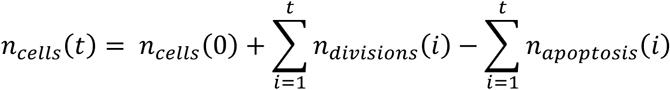

Therefore, the cell fold change within the tissue from *t=* 0 is given by:

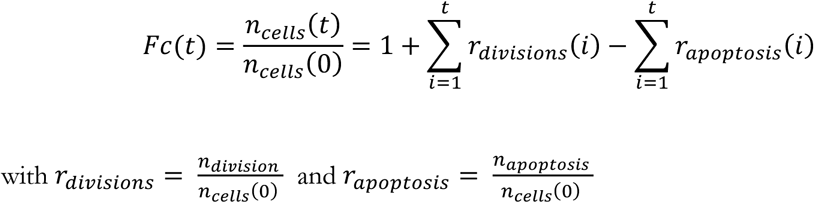

Thus, averaging across animals gives:

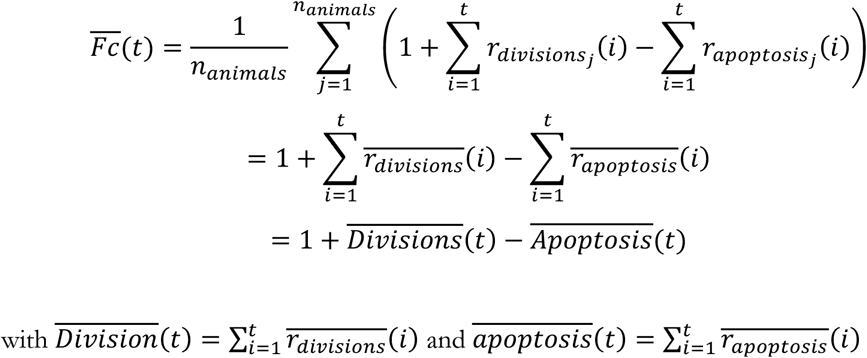

Therefore, allowing to isolate the average contributions of divisions and apoptosis to the average fold change in cell number.

To evaluate the contribution of the compensation through both apoptosis and division rates in *mud* tissue, we considered two different cases:

- The first one, to evaluate the evolution of cell fold change in the absence of compensation by variations in the rate of cell division or apoptosis, 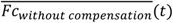. To do so, we evaluated the success rate of division producing two daughter cells 𝑠̅ in *mud* tissue through time as follows:

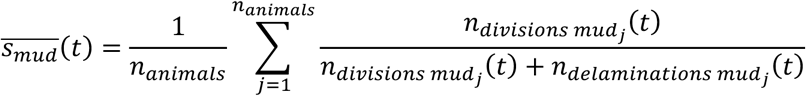

This way, we could compute the adjusted contribution of division to fold change as:

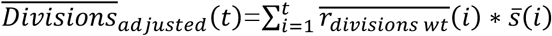

And evaluate the evolution of the fold change as:

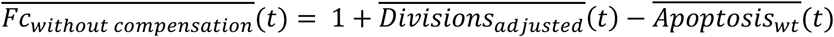

- The second one, to evaluate the contribution of the decreased apoptosis observed in *mud* tissue alone, *Fc_without apoptosis compensation_*. To do so, we replaced the contribution of apoptosis observed in wt to the evolution of the fold change in *mud* tissue as follows:

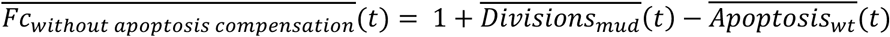

These values were calculated using Python code and plotted to compare the fold change evolution observed in wt and *mud* tissues across different conditions.

#### Quantification of apoptosis beneath the epithelial tissue, and tracking and

Quantification of basal cell apoptosis was performed on maximum intensity Z-stack (80-100 slices at 0.5 µm spacing) projections of fixed tissues acquired using 40x/1.4 OIL DIC H/N2 PL FLUOR or 40x/1.3 OIL DIC H/N2 PL FLUOR objectives. This was achieved by manually counting the number of Casp3-positive corpses underneath the epithelium in a ROI corresponding to a 120.75 µm x 120.75 µm box, using the midline, and the anterior and posterior scutellar macrochaetes as spatial references in tissues counter-stained with antibodies against the SJ proteins Cora or FasIII to reveal the epithelial cells. To evaluate statistical significance between conditions we used either the student *t*-test or ANOVA with post-hoc Tukey test.

To evaluate the fate of delamintated cells in *mud* and *mud grnd^RNAi^*, we manually tracked the cells in tissue expressing the PH:ChFP membrane marker in conjunction with the DNA replication marker PCNA:GFP, which enables to distinguish cells in G1, G2, and S-phase of the cell cycle, as well as mitosis ^95–97^ and apoptosis as evidenced by the fragmentation of the nucleus (Figure S6C). Basally born cells were tracked on average for 4.0 ± 0.2 sem h in Fiji on time-lapse Z-stack live-images (30 slices at 1 µm spacing) aquired using a 100x/1.4 OIL DIC N2 PL APO VC or a 100x/1.45 OIL PL APO L objective at 5 min intervals.

#### 3D cell geometry

To determine the 3D cell geometry in *mud*, *mud pat^RNAi^* and *mud Eb1^RNAi^*, clones expressing a dsRNA against Pat or Eb1 were generated in *mud* tissues expressing the PH:GFP membrane marker. Subsequently, Z-stack live-images were acquired at 0.5 µm spacing using a 100x/1.4 OIL DIC N2 PL APO VC objective. Using the PH:GFP membrane marker we measured the cell height (h) and apical area (A) to determine the 3D cell shape, expressed as 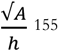, in the double mutant clones and compared it to the *mud* tissue surrounding the double mutant clones. The student *t*-test was used to evaluate statistical significance between cells in *mud* and surrounding double mutant tissue.

#### Quantification of apical protein intensity

Quantification of apical protein intensity (Tub:GFP, Pat:tagRFP and Eb1:tdGFP) during cell reintegration was performed on Z-stack images (20-16 slices at 0.5 µm spacing) acquired at 2 min intervals using a 100x/1.4 OIL DIC N2 PL APO VC or a 100x/1.45 OIL PL APO L objective in tissue also expressing either E-Cad:GFP or E-Cad:3mKate2. To keep the focus of the tissue, autofocus detection (Metamorph software) was used for the duration of the imaging. The apical intensities were measured in Fiji on 2 µm average Z-stack projections around the E-Cad signal and by using the E-Cad signal to delineate the apical areas of the small and large siblings starting at 10 min after anaphase elongation (*t=* 0 min).

To measure the relative apical mean fluorescence intensity (MFI) in the reintegrating cell, and to enable to compare it with wt, the measured intensity of the largest sibling was subtracted from the intensity of the smallest sibling (in *mud* corresponding to the reintegrating cells as characterized by an apex area <10% of its sibling) and the resulting intensity differences for each condition (both wt and mutant) were normalized to the highest measured difference. To evaluate statistically significant differences in apical intensities a student *t*-test was performed at *t=* 10 min after anaphase onset.

The absolute MFI in the reintegrating cell was calculated by dividing the measured intensity in the apex of the small reintegrating cell (after subtraction of the background intensity measured in out-of-focus plane) by the intensity measured in the entire tissue (after subtraction of the background intensity measured in out-of-focus plane). A paired *t*-test was performed to assess whether a statistically significant change in MFI occurred during reintegration, as calculated between the average measured intensities at 10-12 min and 36-38 min.

Apical Sqh:3GFP MFI in cell junctions neighboring a reintegrating cell (+1N) in *mud pat^RNAi^* and *mud w^RNAi^* tissues, was measured on average projected images (1 µm) around the E-Cad:3mKate2 signal at *t=* 20-30 min after anaphase onset, that were acquired at 5 min intervals (20 Z-stack slices at 0.5 µm spacing) using a 100x/1.4 OIL DIC N2 PL APO VC or a 100x/1.45 OIL PL APO L objective. The measured bicellular junctional fluorescent intensity, measured between the vertices of neighboring cells using a line width of 0.65 µm in Fiji, was divided by the total MFI within the tissue. The student *t*-test was performed the determine statistically significant differences.

#### Analyses of apical cap protein localization and formation

To evaluate Pat:tagRFP, Tub:RFP and Eb1:tdGFP localization in the apical cap we imaged at high spatial resolution *mud* tissue co-expressing the PLCγPH:GFP (PH:GFP) or PLCγPH:ChFP (PH:ChFP) membrane markers that specifically bind the phosphoinositide, phosphatidylinositol biphosphate (PIP_2_).^34,156^ Due to photobleaching, we imaged only the apical domain using a 100x/1.4 OIL DIC N2 PL APO VC or a 100x/1.45 OIL PL APO L objective, acquiring Z-stack images (12 slices at 0.5 µm spacing) at 2 min intervals. Representative (selected from >*n*= 5 divisions) x-y and x-z images showing Pat:tagRFP, Tub:RFP and Eb1:tdGFP enrichment at *t=* 12 min after anaphase onset in the apical cap as well as PIP_2_ were generated for reintegrating cells following a highly mis-oriented division using Fiji. To highlight apical cap enrichment in PIP_2_ intensity, a semi-quantitative approach was used to render the image in a color-code reflecting the pixel intensity in the image (fire LUT), thereby highlighting the difference in intensity in the apical cap. To illustrate this PIP_2_ enrichment in full 3D, we imaged PH:GFP in *mud* tissue using a 100x/1.4 OIL DIC N2 PL APO VC objective, acquiring Z-stack images (40 slices at 0.5 µm spacing) at 2 min intervals and generated representative (selected from >*n*= 5 divisions) x-y and x-z images in fire LUT using Fiji at *t=* 12 min after anaphase onset. A similar high resolution imaging protocol and was used to evaluate the apical cap localization of Kst:YFP, Ck:GFP, Shot:YFP, Par-1:GFP, Cad99C:GFP, RhoGEF2:sfGFP, Crb:GFP-A, Jup:GFP, Sas:sfGFP.

To analyze apical cap formation in *mud*, *mud pat^RNAi^* and *mud Eb1^RNAi^*, tissues expressing the PH:GFP membrane marker, with or without the YFP:Asl centrosome marker, were imaged at high spatiotemporal resolution (40-50 slices Z-stack with 0.5 µm spacing acquired every 2 min) using either a 100x/1.45 OIL PL APO L or a 100x/1.4 OIL DIC N2 PL APO VC objective. Cell divisions during which an apical cap was detected and subsequently followed either by an reintegration event or a delaminating daughter cell were mannually scored and used to determine differences amongst genotypes. The chi-square test was used to evaluate statistical significance. Representative divisions followed by a reintegration event or the production of only one epithelial daughter were segmented using Imaris software (Oxford Instruments) to follow their 3D geometry.

#### Quantification of Ban3-GFP reporter and apex cell area

To measure Hippo/YAP activity we used the Ban3-GFP reporter, whose nuclear intensity reflects Hippo/YAP pathway activity^88^, and analyzed daughter cells following division during the second division wave between 17 to 27 hAPF^70,71^ within the previously defined ROI^85^ and which was used to evaluate tissue cell number, division, and apoptosis rates. Thereto, we manually measured in Fiji, by drawing the nuclear outline, the mean Ban3-GFP intensity 60 min post anaphase in the daughter cells, in a single Z-plane of acquired Z-stack images (30 slices at 1.0 µm spacing acquired at 5 or 10 min intervals using a 63x/1.4 OIL DICII PL APO objective) at the center of the nucleus (along the AB tissue axis) in tissue expressing both Ban3-GFP and PH:ChFP to label the cell contours. Dividing cells were visually identified by rounding and apical area expansion. The time of maximum apex area expansion and rounding, prior to anaphase elongation, was set as the 0 min reference. The measured mean nuclear intensity was first corrected for background signal by subtracting the mean background signal (e.g. measured in out-of-focus plane), and subsequently divided by the mean tissue intensity measured in the entire tissue at the same Z-plane as for each individual nucleus. For each daughter cell the corresponding apex area was measured at the most apical plane using the PH:ChFP marker. To compare the intensity in *mud* and wt daughter cells, cells were binned according to their estimated initial mother cells apex area. Thereto, we used the sum of the two wt siblings and the apex area of the lone epithelial *mud* daughter 60 min post-anaphase, which we found correlated well with the initial mother cells apex area at -60 min pre-anaphase (R^2^= 0.84 for *n=* 77 wt cells, and R^2^= 0.63 for *n=* 64 cells *mud*, see Figure S5F). To evaluate statistical significance between wt and *mud* apex area bins we used ANOVA with post-hoc Tukey test.

#### Figure display and movies

All movies and images in the manuscript represent maximum intensity projections and were subjected to denoising, bleach correction and contrast enhancement using, Corel PaintShop Pro, Adobe Photoshop and/or Fiji plugins to improve their visual display.

### Statistics

Apart for the full cell tracking in scutellum tissue of *Sas-4* mutants (*N=* 2), for all analyses samples were taken from at least 3 different animals. The number of cells (*n*) or animals (*N*) analyzed are provided in each figure. The statistical tests used for analyses are indicated in the figure legends and each Methods section. An *f*-test was performed to determine whether the variances amongst conditions differed prior to using the student *t*-test.

### Data availability

All the datasets presented in this manuscript are available from the corresponding authors upon reasonable request.

## Key resources table

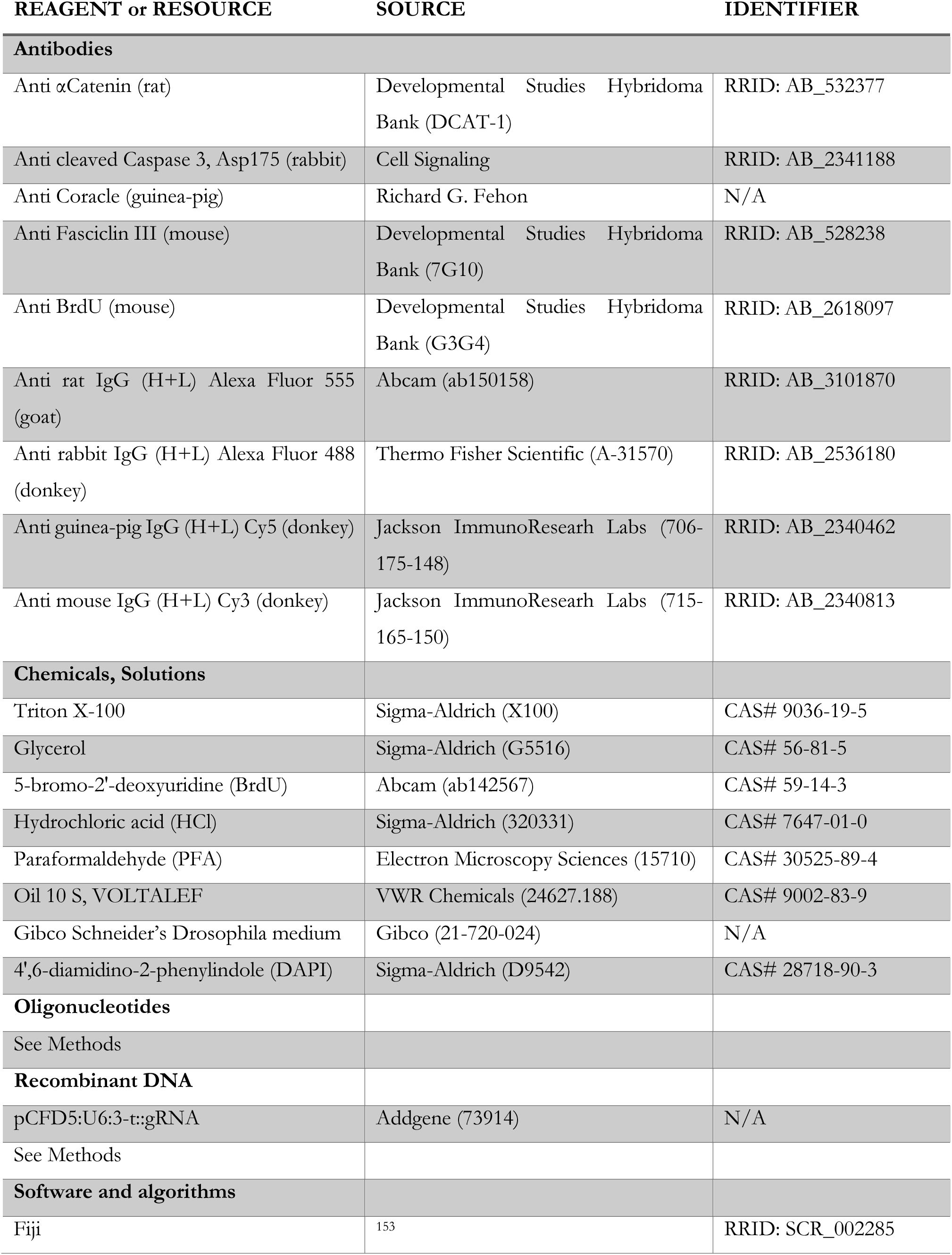
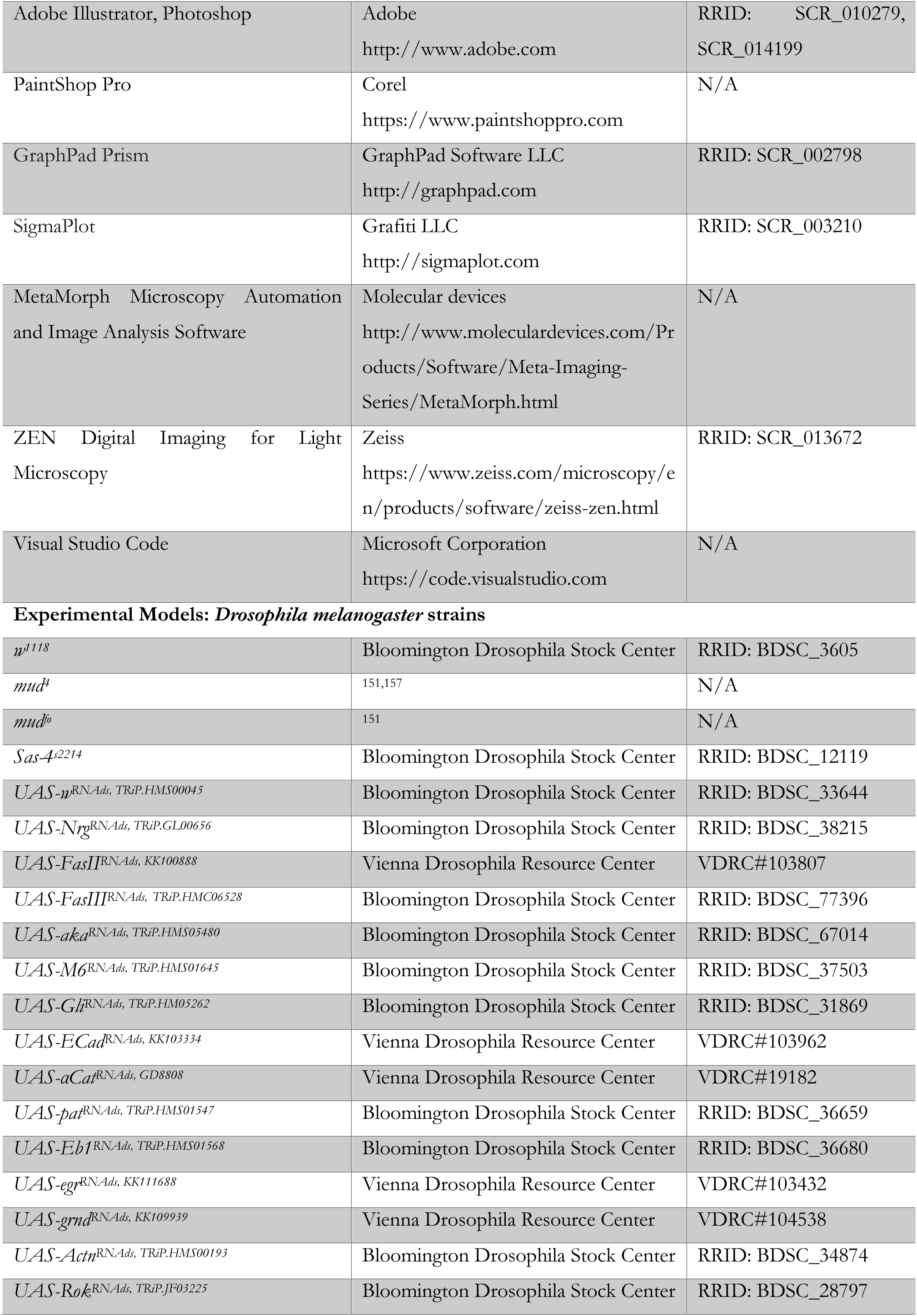
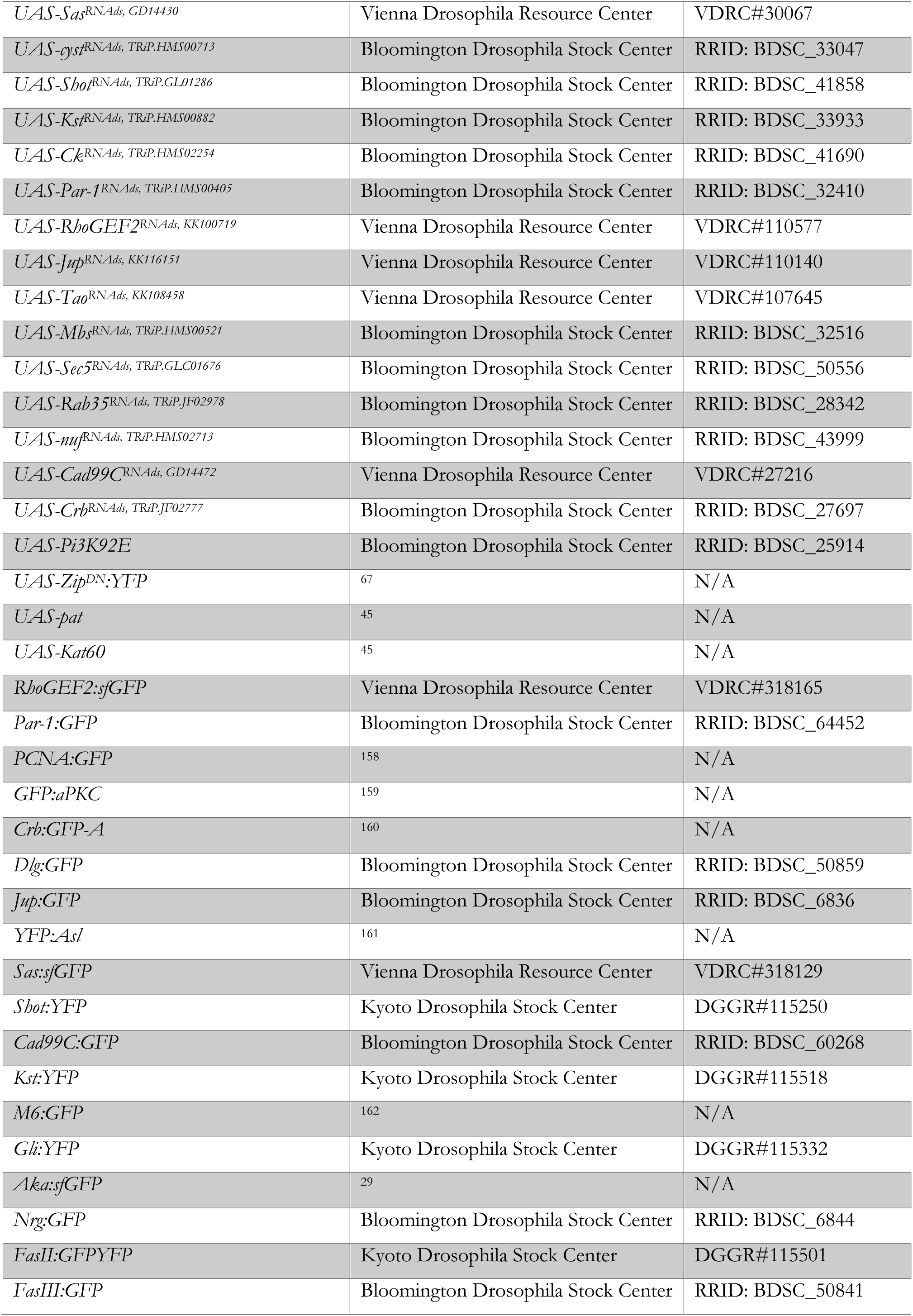
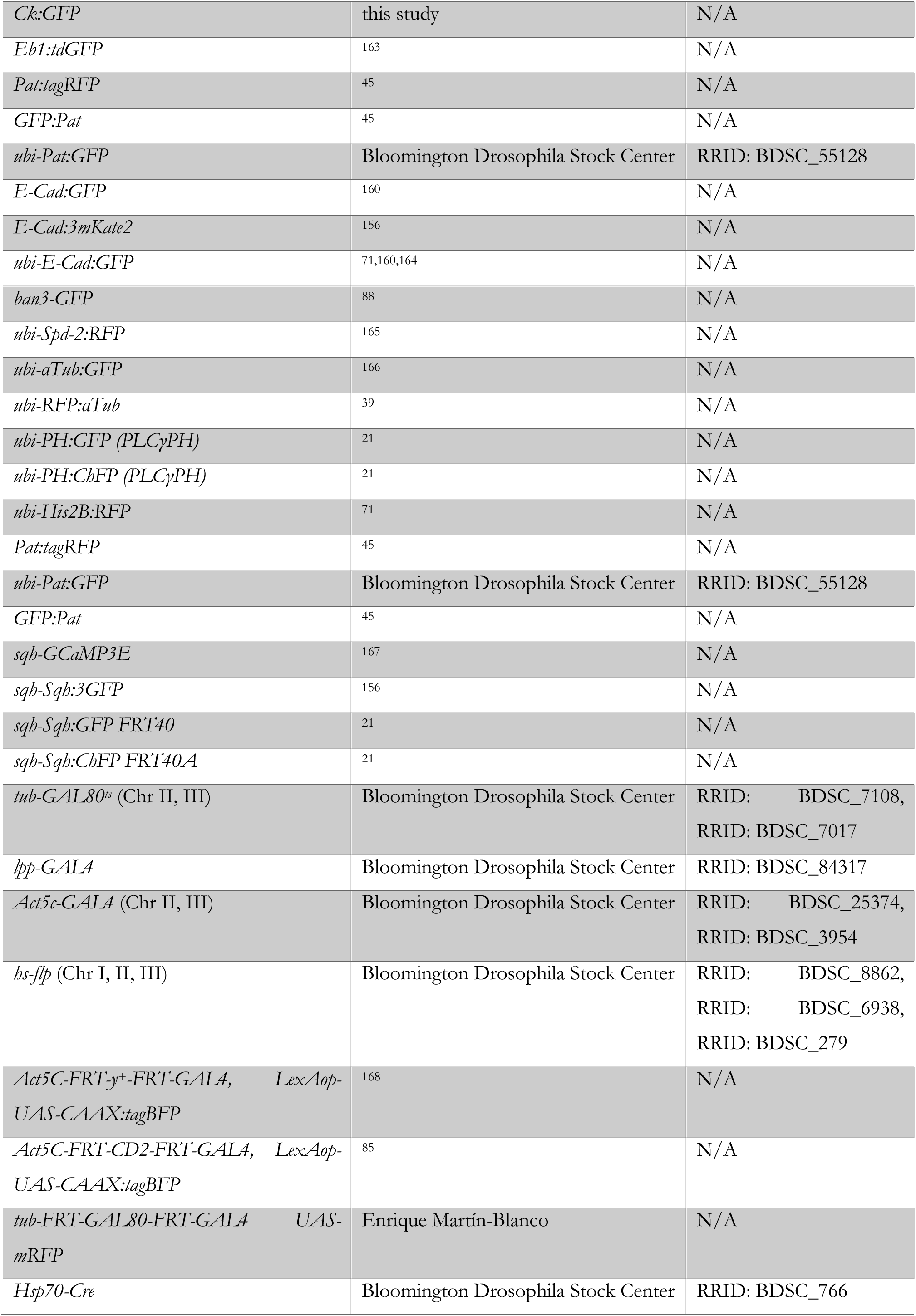
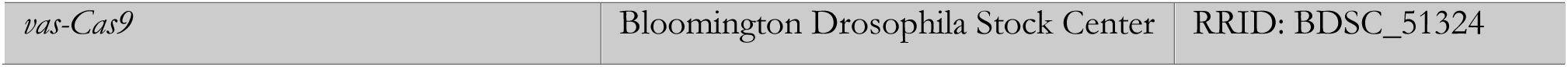

## Acknowledgments

We thank Richard G. Fehon, Stefan Luschnig, Enrique Martín-Blanco, Jose Pastor-Pareja, Yu-Chiun Wang, the Bloomington Drosophila Stock Center, Transgenic RNAi Project at Harvard Medical School, Vienna Drosophila Resource Center, Kyoto Drosophila Stock Center, and the Developmental studies Hybridoma Bank for reagents; the department of genetics and developmental biology imaging facility PICT-IBiSA@BDD for help with microscopy; Lale Alpar, Florencia di Pietro, and Mari W. Yoshida for valuable comments on the manuscript. This work was supported by Institut Curie, CNRS, INSERM, ERC Advanced Scaling-Sensitivity (101020243), ARC (SL220130607097), ANR (TiMecaDiv 20CE13000801), ANR (ChronoDamage 20CE13-0013), CANCERO-INCA (PLBIO2020/BELLAICHE), ANR Labex DEEP (11-LBX-0044, ANR-10-IDEX-0001-02).

## Author contributions

F.B. and Y.B. designed the project and experiments; F.B., E.v.L., S.A., and R.T. performed experiments and analyzed the data; B.T. wrote the code for and ran the simulations; F.B. prepared the figures and videos; Y.B. secured funding for the work; F.B. and Y.B. wrote the manuscript.

---

**Figure S1.**
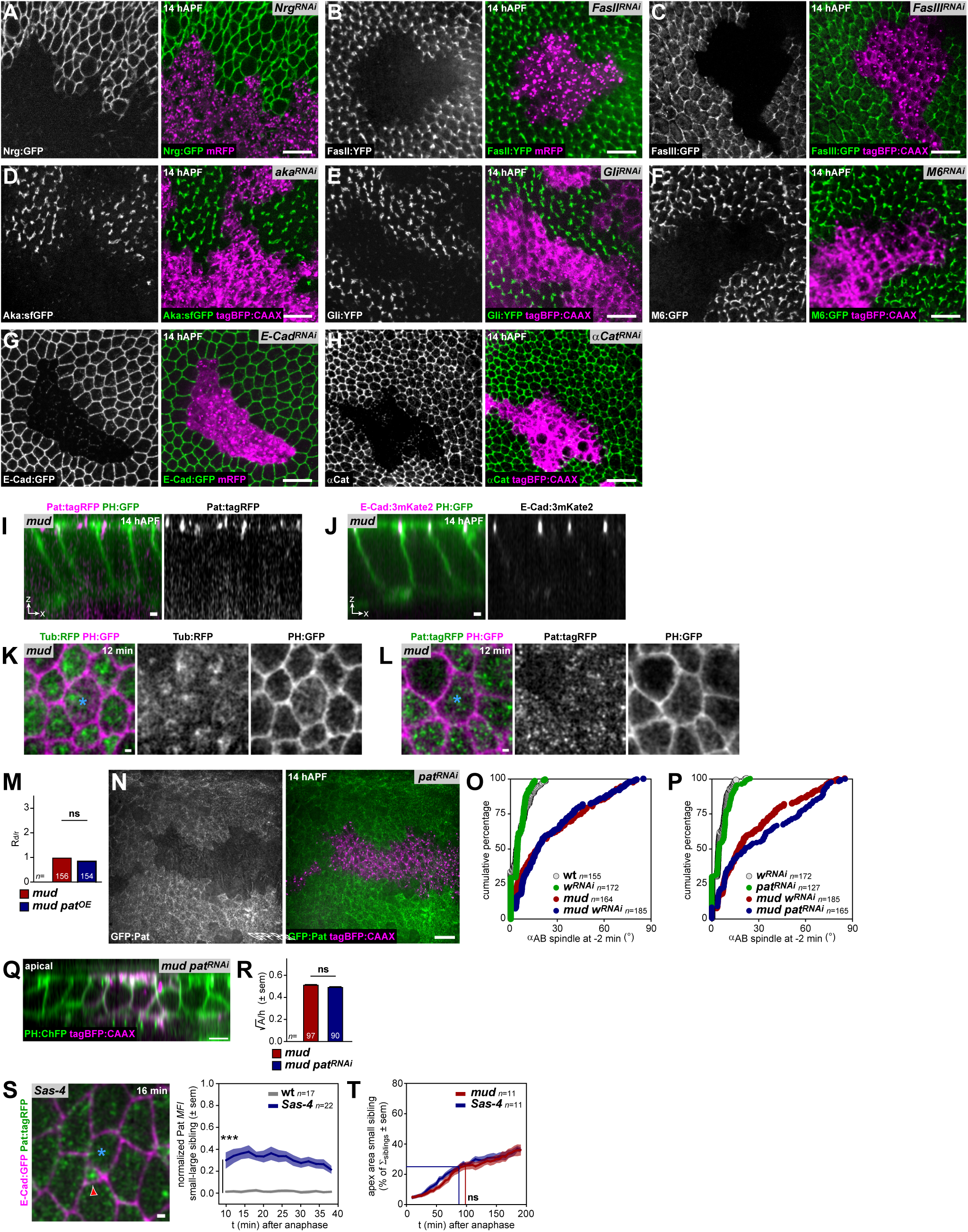
RNAi knock-down efficiency, MT protein localization, intensity and contribution to division orientation, cell shape and reintegration dynamics in wt and mutant conditions. Related to Figure 2. (A) Top view live image of a *Nrg^RNAi^* expressing clone at 14 hAPF (marked by mRFP) in wt notum tissue expressing Nrg:GFP showing the efficacy of Nrg RNAi knock-down. (B) Top view live image of a *FasII^RNAi^* expressing clone at 14 hAPF (marked by mRFP) in wt notum tissue expressing FasII:YFP showing the efficacy of FasII RNAi knock-down. (C) Top view live image of a *FasIII^RNAi^* expressing clone at 14 hAPF (marked by tagBFP:CAAX) in wt notum tissue expressing FasIII:GFP showing the efficacy of FasIII RNAi knock-down. (D) Top view live image of an *aka^RNAi^* expressing clone at 14 hAPF (marked by tagBFP:CAAX) in wt notum tissue expressing Aka:sfGFP showing the efficacy of Aka RNAi knock-down. (E) Top view live image of a *Gli^RNAi^* expressing clone at 14 hAPF (marked by tagBFP:CAAX) in wt notum tissue expressing Gli:YFP showing the efficacy of Gli RNAi knock-down. (F) Top view live image of a *M6^RNAi^* expressing clone at 14 hAPF (marked by tagBFP:CAAX) in wt notum tissue expressing M6:GFP showing the efficacy of M6 RNAi knock-down. (G) Top view live image of an *E-Cad^RNAi^* expressing clone at 14 hAPF (marked by mRFP) in wt notum tissue expressing E-Cad:GFP showing the efficacy of E-Cad RNAi knock-down. (H) Top view image of a á*Cat^RNAi^* expressing clone at 14 hAPF (marked by tagBFP:CAAX) in a fixed wt notum tissue labelled with antibodies against αCat showing the efficacy of αCat RNAi knock-down. (I) AB view live image showing apical Pat:tagRFP localization in *mud* tissue co-expressing PH:GFP. (J) AB view live image showing apical E-Cad:3mKate2 localization in *mud* tissue co-expressing PH:GFP. (K) Top view live images of apical Tub:RFP and PH:GFP localization in a lone daughter cell (blue asterisk) at *t*= 12 min after anaphase onset following a mis-oriented division producing a single epithelial daughter and a daughter that failed to reintegrate (note the absence of an apical cap). Representative images of *n=* 9 divisions. (L) Top view live images of apical Pat:tagRFP and PH:GFP localization in a lone daughter cell (blue asterisk) at *t*= 12 min after anaphase onset following a mis-oriented division producing a single epithelial daughter and a daughter that failed to reintegrate (note the absence of an apical cap). Representative images of *n=* 8 divisions. (M) Graph of the delamination/reintegration ratio (R_d/r_) in *mud*, and *mud pat^OE^* tissue normalized to the R_d/r_ in *mud*. ns, not significant, chi-square test. *n*, number of divisions analyzed. (N) Top view live image of a *pat^RNAi^* expressing clone at 14 hAPF (marked by tagBFP:CAAX) in wt notum tissue expressing Pat:GFP showing the efficacy of Pat RNAi knock-down. (O) Graph of áAB at *t=* -2 min prior to anaphase onset in wt, *wRNAi*, *mud* and *mud w^RNAi^*, plotted as cumulative percentage. The αAB in wt and *wRNAi* are not different (*p*<0.8690, Mann-Whitney U test). The αAB in *mud* and *mud wRNAi* are not different (*p*<0.4277, Mann-Whitney U test). *n*, number of cells analyzed. (P) Graph of the AB spindle orientation (áAB) at *t=* -2 min prior to anaphase onset in *wRNAi*, *patRNAi*, *mud w^RNAi^* and *mud pat^RNAi^*, plotted as cumulative percentage. The αAB in *wRNAi* and *patRNAi* are not different (*p*<0.1192, Mann-Whitney U test). The αAB in *mud w^RNAi^* and *mud pat^RNAi^* are not different (*p*<0.1778, Mann-Whitney U test). *n*, number of cells analyzed. (Q) AB view live image of *mud* tissue expressing the PH:ChFP membrane marker and harboring a *pat^RNAi^* expressing clone at 14 hAPF (marked by tagBFP:CAAX). (R) Graph of the 3D cell geometry measured as the square-root of the cell apex area divided by the cell height (mean ± sem) in *mud* and neighboring *mud pat^RNAi^* mosaic tissue. ns, not significant, student *t*-test. *n*, number of cells analyzed. (S) The left top view live image shows apical Pat:tagRFP localization at *t=* 16 min after anaphase onset in a small daughter cell (red arrowhead) following a highly mis-oriented division in *Sas-4* tissue co-expressing E-Cad:GFP. Blue asterisk, large sibling. The right graph shows the evolution of the normalized Pat:tagRFP mean apical fluorescence intensity (*MFI* ± sem) measured in the smallest daughter cells as compared to their larger siblings upon division in wt and *Sas-4* tissues. *** *p*<0.0001, student *t*-test at *t=* 10 min. *n*, number of cells analyzed. (T) Graph of the apex area dynamics of the small daughter cells (initial apex area <10% of its sibling) in *mud* and *Sas-4* tissues (mean % of the sum of the apex areas are of the two siblings ± sem, sliding average 30 min). Lines indicate t_1/2_, the time to reach halfway of the predicted area (i.e. 50% of the mother apex area). The t_1/2_ of mispositioned cells is not significantly different between *mud* and *Sas-4* tissues. ns, not significant, student *t*-test. *n*, number of cells analyzed. Scale bars: 1 µm (I-L, S), 5 µm (Q), 15 µm (A-H, N).

**Figure S2.**
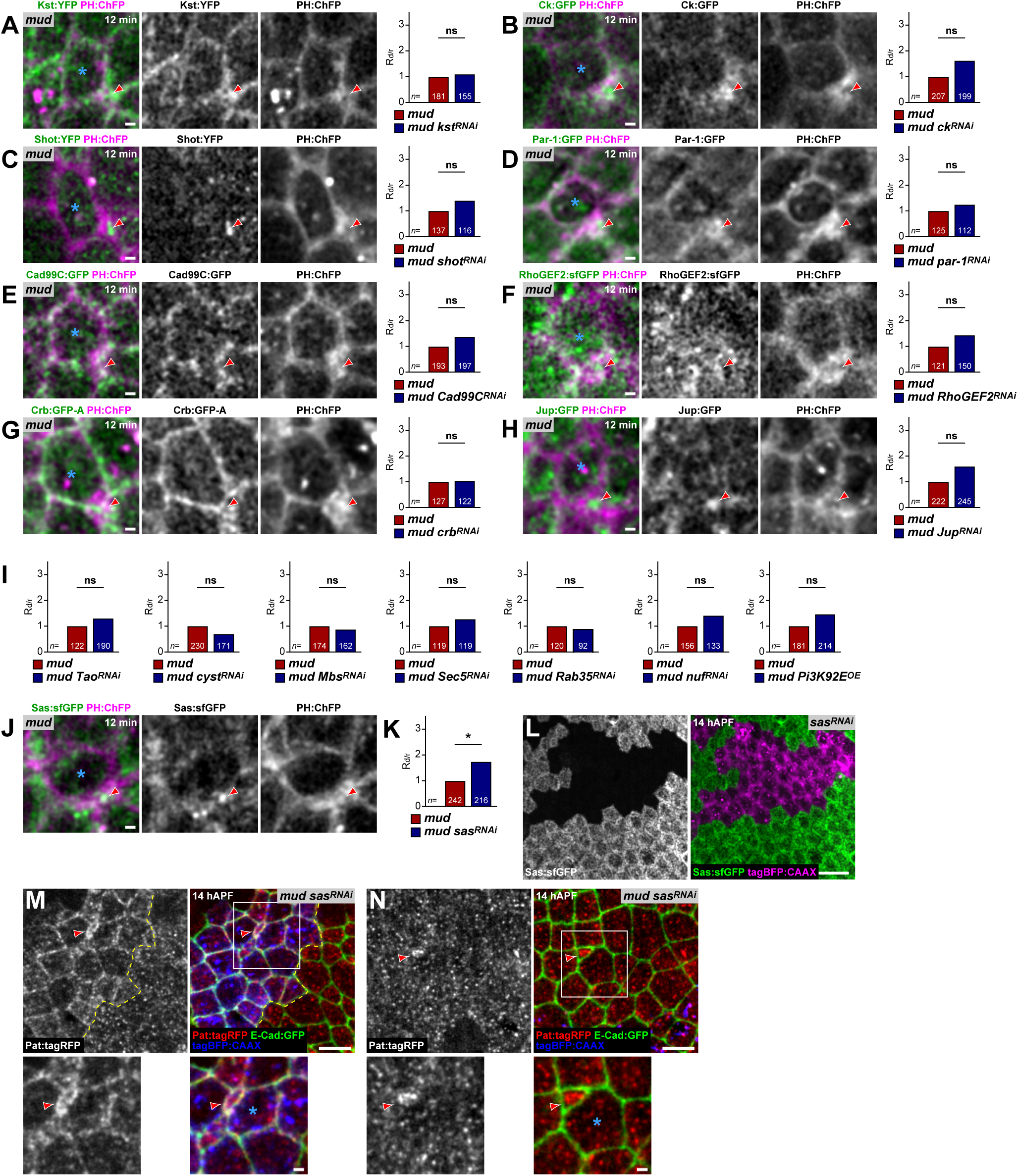
The apical transmembrane receptor Sas modulates Pat localization and cell reintegration. Related to Figure 3. (A-H) Top view live images of the apical cap (red arrowheads) localization of selected protein candidates at *t=* 12 min after anaphase onset upon cell reintegration in *mud* tissue co-expressing PH:ChFP: Kst:GFP (A), Ck:GFP (B), Shot:YFP (C), Par-1:GFP (D), Cad99C:GFP (E), RhoGEF2:sfGFP (F), Crb:GFP-A (G), Jup:GFP (H). Representative images of *n= >*5 divisions. Blue asterisk, large sibling. For each condition, the graph of the delamination/reintegration ratio (R_d/r_) in *mud*, and double mutant tissue normalized to the R_d/r_ in *mud* is shown (right graph of each panel). ns, not significant, chi-square test. *n*, number of divisions analyzed. (I) Graphs of the delamination/reintegration ratios (R_d/r_) in *mud*, and *mud Tao^RNAi^*, *mud cyst^RNAi^*, *mud Mbs^RNAi^*, *mud Sec5^RNAi^*, *mud Rab35^RNAi^*, *mud nuf^RNAi^*, *mud Pi3K92E^OE^* tissue normalized to the R_d/r_ in *mud*. ns, not significant, chi-square test. *n*, number of divisions analyzed. (J) Top view live images of the apical cap (red arrowheads) localization of Sas:sfGFP and PH:ChFP at *t=* 12 min after anaphase onset upon cell reintegration in *mud* tissue. Representative images of *n= >*5 divisions. Blue asterisk, large sibling. (K) Graph of the delamination/reintegration ratio (R_d/r_) in *mud*, and *mud sas^RNAi^* tissue normalized to the R_d/r_ in *mud*. * *p*<0.05, chi-square test. *n*, number of divisions analyzed. (L) Top view live image of a *sas^RNAi^* expressing clone at 14 hAPF (marked by tagBFP:CAAX) in wt notum tissue expressing Sas:sfGFP showing the efficacy of Sas RNAi knock-down. (M-N) Top view live images of a *sas^RNAi^* clone (marked by tagBFP:CAAX, indicated by the yellow dashed lines) in *mud* tissue at 14 hAPF showing Pat:tagRFP and E-Cad:GFP localization in a reintegrating cell (red arrowheads) in double mutant *mud sas^RNAi^* (M) and surrounding *mud* tissue (N). Note the apical Pat:tagRFP delocalization from the medial cell apex to the cell junctions upon *sas^RNAi^*. Bottom images are close-ups of the boxed regions in M and N, respectively (scale bars, 1 µm). Blue asterisks, large siblings. Scale bars: 1 µm (A-H, J), 10 µm (M, N), 15 µm (L).

**Figure S3.**
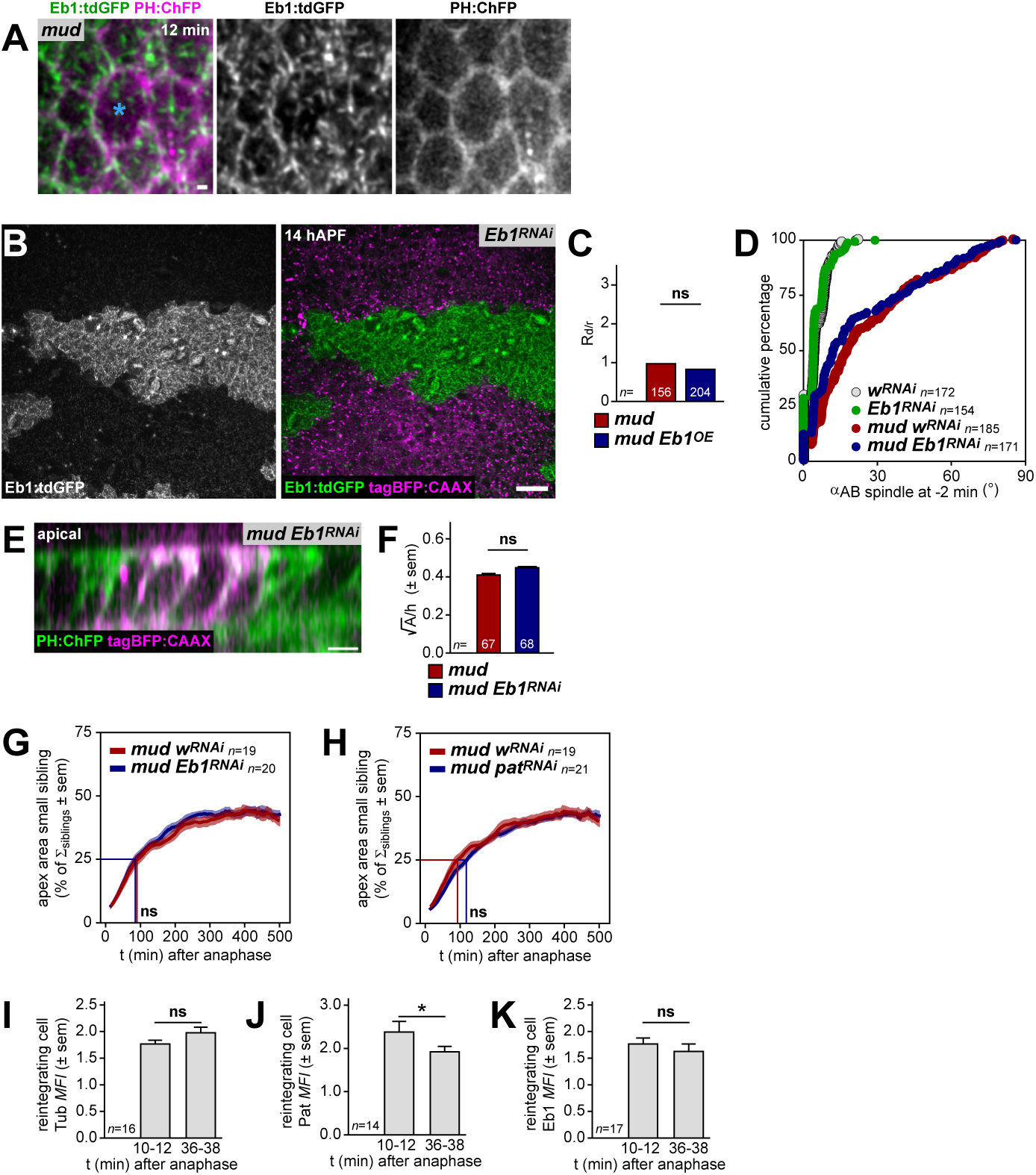
Eb1 localization, contribution to division orientation, cell shape and reintegration dynamics in wt and *mud*. Related to Figure 3. (A) Top view live images of apical Eb1:tdGFP and PH:ChFP localization in a lone daughter cell (blue asterisk) at *t*= 12 min after anaphase onset following a mis-oriented division producing a single epithelial daughter and a daughter that failed to reintegrate (note the absence of an apical cap). Representative images of *n=* 13 divisions. (B) Top view live image of a *Eb1^RNAi^* expressing clone at 14 hAPF (marked by tagBFP:CAAX) in wt notum tissue expressing Eb1:tdGFP showing the efficacy of Eb1 RNAi knock-down. (C) Graph of the delamination/reintegration ratio (R_d/r_) in *mud*, and *mud Eb1^OE^* tissue normalized to the R_d/r_ in *mud*. ns, not significant, chi-square test. *n*, number of divisions analyzed. (D) Graph of áAB at *t=* -2 min prior to anaphase onset in *w^RNAi^*, *Eb1^RNAi^*, *mud w^RNAi^* and *mud Eb1^RNAi^*, plotted as cumulative percentage. The αAB in *w^RNAi^* and *Eb1^RNAi^* are not different (*p*<0.5512, Mann-Whitney U test). The αAB in *mud w^RNAi^* and *mud Eb1^RNAi^* are not different (*p*<0.0775, Mann-Whitney U test). *n*, number of cells analyzed. (E) AB view live image of *mud* tissue expressing the PH:ChFP membrane marker and harboring an *Eb1^RNAi^* expressing clone at 14 hAPF (marked by tagBFP:CAAX). (F) Graph of the 3D cell geometry measured as the square-root of the cell apex area divided by the cell height (mean ± sem) in *mud* and neighboring *mud Eb1^RNAi^* mosaic tissue. ns, not significant, student *t*-test. *n*, number of cells analyzed. (G) Graph of the apex area dynamics of the small daughter cells (initial apex area <10% of its sibling) in *mud w^RNAi^* and *mud Eb1^RNAi^* tissues (% of the sum of the apex areas of the two siblings ± sem, sliding average 30 min). Lines indicate t_1/2_, the time to reach halfway of the predicted area (i.e. 50% of the mother apex area). The t_1/2_ of mispositioned cells is not significantly different between *w^RNAi^* and *mud Eb1^RNAi^* tissues. ns, not significant, student *t*-test. *n*, number of cells analyzed. (H) Graph of the apex area dynamics of the small daughter cells (initial apex area <10% of its sibling) in *mud w^RNAi^* and *mud pat^RNAi^* tissues (mean % of the sum of the apex areas of the two siblings ± sem, sliding average 30 min). Lines indicate t_1/2_, the time to reach halfway of the predicted area (i.e. 50% of the mother apex area). The t_1/2_ of mispositioned cells is not significantly different between *w^RNAi^* and *mud pat^RNAi^* tissues. ns, not significant, student *t*-test. *n*, number of cells analyzed. (I) Graph of the Tub:GFP mean apical fluorescence intensity (*MFI* ± sem) in reintegrating cells in *mud* tissue at two time points after anaphase onset (MFI at 10-12 min and 36-38 min were averaged). ns, not significant, student *t*-test. *n*, number of cells analyzed (for the same cells the evolution of the apical MFI in the small sibling as compared to its larger daughters are shown in Figure 2H). (J) Graph of the Pat:tagRFP mean apical fluorescence intensity (*MFI* ± sem) in reintegrating cells in *mud* tissue at two time points after anaphase onset (MFI at 10-12 min and 36-38 min were averaged). * *p*<0.05, student *t*-test. *n*, number of cells analyzed (for the same cells the evolution of the apical MFI in the small sibling as compared to its larger daughters are shown in Figure 3B). (K) Graph of the Eb1:tdGFP mean apical fluorescence intensity (*MFI* ± sem) in reintegrating cells in *mud* tissue at two time points after anaphase onset (MFI at 10-12 min and 36-38 min were averaged). ns, not significant, student *t*-test. *n*, number of cells analyzed (for the same cells the evolution of the apical MFI in the small sibling as compared to its larger daughters are shown in Figure 3E). Scale bars: 1 µm (A), 5 µm (E), 15 µm (C).

**Figure S4.**
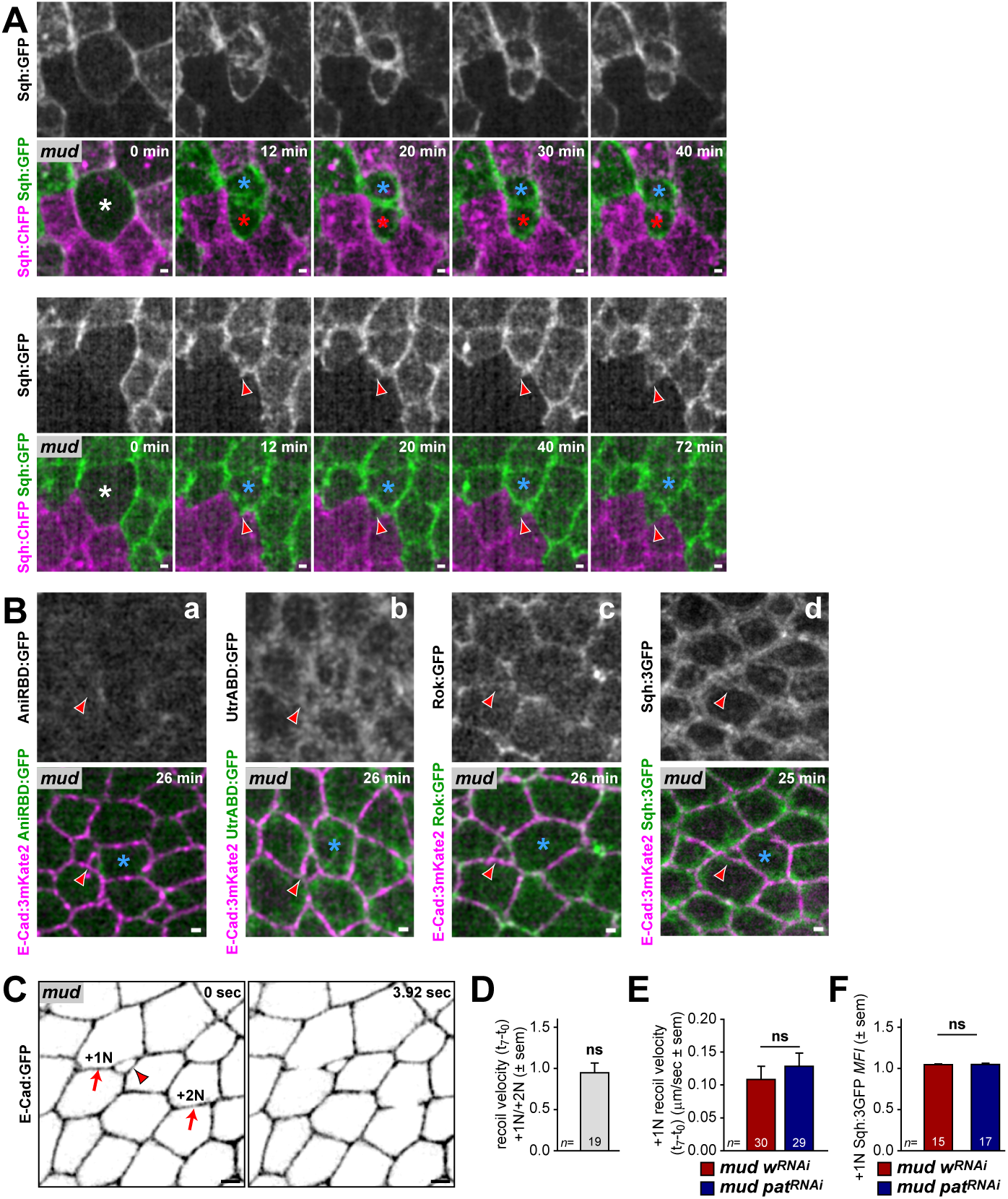
Apical actomyosin localization and analyses of junction tension during cell reintegration in *mud* and upon Pat loss of function. Related to Figure 4. (A) Top view time-lapse images of apical MyoII regulatory light chain Sqh:GFP and Sqh:ChFP ^152^patches^169^ in *mud* tissue following a planar cell division (top panels) and a highly mis-oriented division producing a reintegrating daughter cell (bottom panels, red arrowheads). Representative images of *n=* 5 divisions. White asterisks, dividing cell at anaphase onset *t*= 0 min. Blue and red asterisks indicate tissue resident daughter cells following division. (B) Top view live images at *t=* 25-26 min after anaphase onset of apical AniRBD:GFP (a), UtrABD:GFP (b), Rok:GFP (c), and Sqh:3GFP (d) as well as E-Cad:3mKate2 in *mud* tissue following a highly mis-oriented division producing a reintegrating daughter cell (red arrowheads). Representative images of *n= >*3 divisions. Blue asterisks, tissue resident daughter cells following division. (C) Top view time-laps images of apical E-Cad:GFP in *mud* tissue prior to (*t_0_*= 0 sec) and after ablation (*t_7_*= 3.92 sec) of a cell junction neighboring a reintegrating cell (+1N) as well as a junction belonging to a cell 2 cell distances away from a reintegrating cell (+2N). Red arrowhead, reintegrating cell. Red arrows, sites of junction ablation. (D) Graph of the +1N/+2N ratio of the recoil velocity (between *t_7_*-*t_0_*, mean ± sem) upon laser ablation of cell junctions neighboring a reintegrating cell (+1N) as well as junctions belonging to a cell 2 cell distances away from the reintegrating cell (+2N) in *mud* tissue. ns, not significant, paired student *t*-test. *n*, number of cells analyzed. (E) Graph of the +1N recoil velocity (between *t_7_*-*t_0_*, mean ± sem) upon laser ablation of cell junctions neighboring a reintegrating cell in *mud w^RNAi^* and *mud pat^RNAi^* tissue. ns, not significant, student *t*-test. *n*, number of cells analyzed. (F) Graph of the +1N Sqh:3GFP mean apical fluorescence intensity (*MFI* ± sem) at cell junctions neighboring a reintegrating cell in *mud w^RNAi^* and *mud pat^RNAi^* tissue. ns, not significant, student *t*-test. *n*, number of cells analyzed. Scale bars: 1 µm (A, B), 2 µm (C).

**Figure S5.**
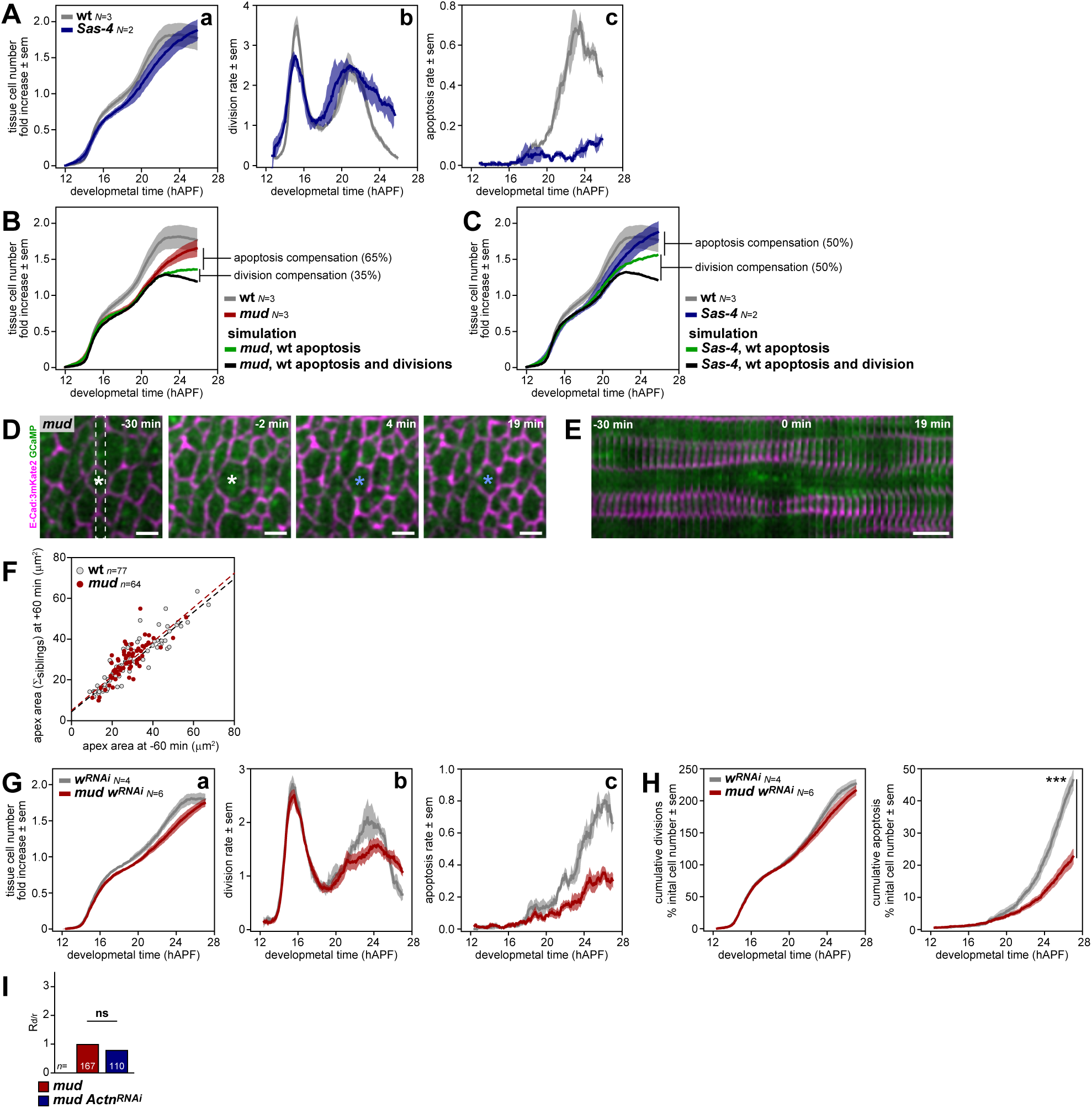
Characterization of wt, *mud* and *Sas-4* cell and tissue dynamics. Related to Figure 5. (A) Graphs of the tissue cell number fold increase (a)(relative to the initial cell number, mean ± sem), the total division rate (b), and the apoptosis rate (c)(rates relative to the initial cell number, mean ± sem, sliding average 60 min) in wt and *Sas-4* scutellum tissue. *N*, hemi-scutella analyzed. (B) Graph of the tissue cell number fold increase (relative to the initial cell number, mean ± sem) in wt and *mud* scutellum tissue, as well as the simulated tissue cell number fold increase in *mud* with the wt apoptosis rate (green curve), and *mud* without compensatory mechanisms, hence, incorporating the wt apoptosis and division rates (blue curve). Note that for the simulation of *mud* without compensatory mechanisms, the rate of successful divisions producing two daughter cells in *mud* is applied to the wt division rate (see Methods). The simulation curves illustrate the extent to which apoptosis 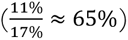 and division 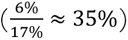 compensate for the cell loss due to mis-oriented divisions. *N*, hemi-scutella analyzed. For wt and *mud* the same data are shown in Figure 5A. (C) Graph of the tissue cell number fold increase (relative to the initial cell number, mean ± sem) in wt and *Sas-4* scutellum tissue, as well as the simulated tissue cell number fold increase in *Sas-4* with the wt apoptosis rate (green curve), and *Sas-4* without compensatory mechanisms, hence, incorporating the wt apoptosis and division rates (blue curve). Note that for the simulation of *Sas-4* without compensatory mechanisms, the rate of successful divisions producing two daughter cells in *Sas-4* is applied to the wt division rate (see Methods). The simulation curves illustrate the extent to which apoptosis 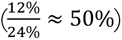 and division 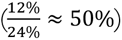 compensate for the cell loss due to mis-oriented divisions. *N*, hemi-scutella analyzed. For wt and *Sas-4* the same data are shown in Figure S5A. (D, E) Representative time-lapse images (D) and kymograph (E) of a highly mis-oriented division producing a lone epithelial daughter in *mud* tissue expressing E-Cad:3mKate2 and GCaMP Ca^2+^ reporter. Note that mis-oriented division is not associated with elevated calcium signaling. Representative images of *n=* 7 divisions. *t=* 0 min, anaphase onset. Dashed box, region plotted as a kymograph. White asterisks, dividing cell. Blue asterisks, lone sibling. (F) Graph of the correlation between the apex area of the mother cell -60 min prior to anaphase onset (*t=* 0 min) and the area of the daughter cells (sum of the area of the siblings for wt and the area of the lone sibling in *mud* after division mis-orientation) +60 min after anaphase onset. Both correlate well in wt and *mud* tissues (dashed lines). R^2^ for wt (0.84) and *mud* (0.63). *n*, number of divisions analyzed. (G) Graphs of the tissue cell number fold increase (a)(relative to the initial cell number, mean ± sem), the total division rate (b), and the apoptosis rate (c)(rates relative to the initial cell number, mean ± sem, sliding average 60 min) in wt *w^RNAi^* and *mud w^RNAi^* medial scutellum tissue. *N*, hemi-scutella analyzed. (H) Graphs of the cumulative divisions and apoptosis (% of initial cell number, mean ± sem) in wt *w^RNAi^* and *mud w^RNAi^* medial scutellum tissue. The final cumulative apoptosis is significantly different between wt *w^RNAi^* and *mud w^RNAi^*. *** *p*<0.001, student *t*-test. *N*, hemi-scutella analyzed. (I) Graph of the delamination/reintegration ratio (R_d/r_) in *mud* and *mud Actn^RNAi^* tissue normalized to the R_d/r_ in *mud*. ns, not significant, chi-square test *n*, number of divisions. Scale bars: 5 µm (D), 10 min (E).

**Figure S6.**
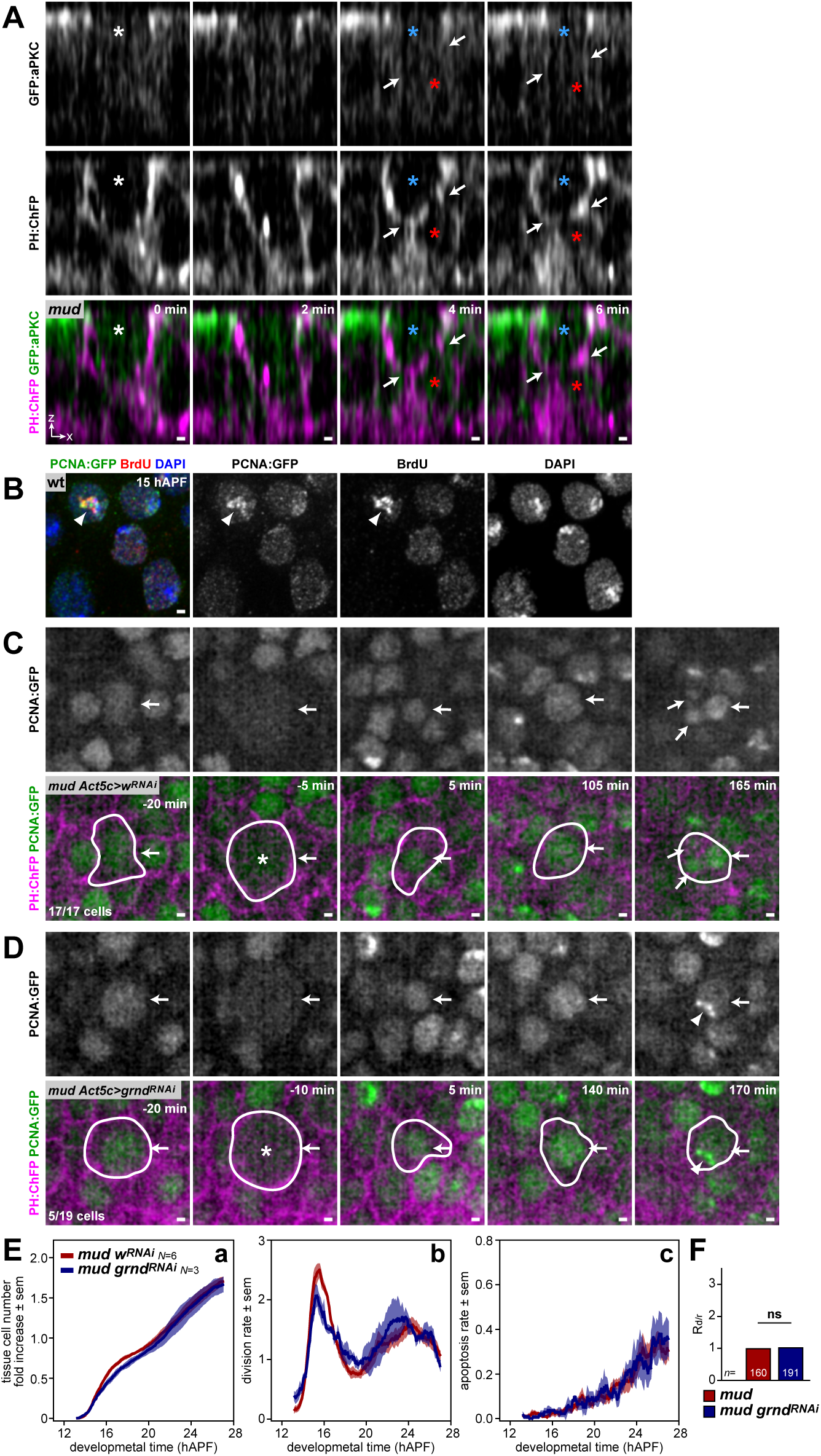
Basally born cells destined to delaminate lack aPKC and can renter S-phase upon *grnd^RNAi^*, but do not contribute to tissue cell number regulation. Related to Figure 6. (A) AB view time-lapse images of PH:ChFP and GFP:aPKC in *mud* tissue upon a highly mis-oriented division producing a lone epithelial daughter cell. The apical GFP:aPKC marker does not accumulate at the most apical domain (white arrows) in the basally born daughter cell (red asterisks). Representative images of *n=* 5 divisions. White asterisk, dividing cell at anaphase onset *t*= 0 min. Blue asterisks, tissue remaining sibling. (B) Fixed wt notum tissue top view image showing PCNA:GFP and BrdU colocalization at 15 hAPF after a 30 min pulse of BrdU incorporation in *ex vivo* culture. Note that PCNA:GFP replication foci colocalize with sites of strong BrDU incorporation (white arrowheads). DNA is labelled with DAPI. (C) Top view time-lapse images of PCNA:GFP and PH:ChFP from -20 min to 165 min after anaphase onset (*t=* 0 min) of a dividing cell in *mud w^RNAi^* tissue producing a lone epithelial daughter (white arrows). In time the delaminated cell undergoes nucleus fragmentation as evidenced by the presence of multiple PCNA:GFP structures within the cell (*t=* 165 min). 17 out of 17 tracked cell displayed such apoptotic feature. White asterisk, dividing cell. White outlines, cell mask based on PH:ChFP labelling. (D) Top view time-lapse images of PCNA:GFP and PH:ChFP from -20 min to 170 min after anaphase onset (*t=* 0 min) of a dividing cell in *mud grnd^RNAi^* tissue producing a lone epithelial daughter (white arrows). In time the delaminated cell undergoes S-phase entry as evidenced by the presence of multiple PCNA:GFP DNA replication foci within the nucleus (white arrowhead at *t=* 170 min). 5 out of 19 tracked cell displayed such reentry into S-phase. White asterisk, dividing cell. White outlines, cell mask based on PH:ChFP labelling. (E) Graphs of the tissue cell number fold increase (a)(relative to the initial cell number, mean ± sem), the total division rate (b) and the apoptosis rate (c)(rates relative to the initial cell number, mean ± sem, sliding average 60 min) in *mud Act5c>w^RNAi^* and *mud Act5c>grnd^RNAi^* tissues. *N*, hemi-scutella analyzed. (F) Graph of the delamination/reintegration ratio (R_d/r_) in *mud* and *mud grnd^RNAi^* tissue normalized to the R_d/r_ in *mud*. ns, not significant, chi-square test. *n*, number of divisions analyzed. Scale bars: 1 µm.

**Video S1.**

Apical top view time-lapse series of dividing cells (asterisks) in wt (left) and *mud* notum tissue expressing E-Cad:GFP. The second panel shows a *mud* division producing two daughter cells of unequal apex area, the third panel a highly mis-oriented division resulting in a mispositioned daughter cell that reintegrates back within the epithelium (arrows), while the right panel shows a division that result in the production of only one epithelial daughter cell.

**Video S2.**

3D segmented time-lapse series of a highly mis-oriented division in *mud* tissue showing the reintegration of a mispositioned daughter cell (red cell) relative to its sibling with larger apex area (blue cell) (left), as well as a mis-oriented division producing a lone epithelial daughter (blue cell) and a delaminating daughter (red cell) (right).

**Video S3.**

Apical top view time-lapse series of Tub:GFP and E-Cad:3mKate2 (top panels), Pat:tagRFP and E-Cad:GFP (middle panels), and Eb1:GFP and E-Cad:3mKate2 (bottom panels) in *mud* tissue showing a highly mis-oriented division (white asterisk) resulting in the reintegration of a mispositioned daughter (arrows). Note that Tub:GFP, Pat:tagRFP and Eb1:GFP are enriched in the smaller sibling as compared to its larger sibling.

**Video S4.**

Apical top view time-lapse series of tissue expressing the PH:GFP membrane marker illustrating the formation of an apical cap preceding cell reintegration (arrows) and absence of apical cap formation upon division that resulted in the formation of only one epithelial daughter in *mud* tissue (top left and right respectively). Initiation of apical cap formation (arrows) is observed in *mud pat^RNAi^* and *mud Eb1^RNAi^* tissues, but is followed by the production of only one epithelial daughter cell (bottom left and right respectively). Asterisks, dividing cells.

**Video S5.**

Apical top view time-lapse series of Jup:GFP and E-Cad:3mKate2 in *mud* tissue following ablation of the medial cortex and of the surrounding cells (top versus bottom panels) of a mispositioned daughter with small apex area (white asterisks).

**Video S6.**

Apical top view time-lapse series of E-Cad:3mKate2 showing a reintegrating daughter cell (arrows) following a highly mis-oriented division (asterisk) in *mud zip^DN^* tissue (left), and a reintegrating daughter cell (arrows) upon a highly mis-oriented division (asterisk) in *mud* tissue neighboring *mud* tissue over-expressing *zip^DN^* (indicated by yellow dashed lines)(right).

